# Isotopic tracing in *Mycobacterium tuberculosis*-infected alveolar epithelial cells indicates metabolic specificity to their virulence and drug-resistance status

**DOI:** 10.1101/2024.12.03.625217

**Authors:** Subia Akram, Shyam Kumar Masakapalli, Ranjan Kumar Nanda

## Abstract

*Mycobacterium tuberculosis* (Mtb), the causative organism of tuberculosis (TB), infects different host cells; however, studies on its metabolic impact on the type 2 alveolar epithelial cells are limited. In this study, comparative metabolic insights of A549 cells infected with laboratory (H37RV, H37Ra) and clinical isolates (drug-resistant: S6, S11 and sensitive: S4, S5) were derived from read out of the ^13^C-based proteinogenic amino acid kinetics. Mtb H37Rv (virulent) exhibited higher growth kinetics within A549 cells compared to H37Ra (avirulent), whereas drug-resistant clinical isolate S6 showed the highest intracellular growth. The viability of H37Rv-infected A549 cells was significantly low. Interestingly, the drug-resistant clinical Mtb isolates maintained better cell viability over time. A [^13^C_6_] glucose tracer and mass isotopomer distributions revealed that A549 cells infected with virulent Mtb strains exhibited higher dependency on the central carbon metabolism like glycolysis, the pentose phosphate pathway and the tricarboxylic acid cycle for *de novo* amino acid biosynthesis. Drug-resistant Mtb isolates infected A549 cells showed robust ^13^C incorporation in the proteinogenic Threonine, Methionine and Lysine, derived from the tricarboxylic acid cycle. A549 cells infected with Mtb, irrespective of their drug resistance status, showed ^13^C incorporation in the essential amino acids (lysine, methionine, threonine, valine), underlining the dependencies of the pathogen on the *de novo* host metabolic precursors. These findings elucidate the importance of deciphering the strain-specific host-pathogen metabolic interactions to develop targeted therapies against drug-resistant Mtb infections.

## Introduction

*Mycobacterium tuberculosis* (Mtb), an intracellular pathogen, infects the lung cells after inhalation.^1^ Lung macrophages phagocytise Mtb and interact with other immune cells like dendritic cells (DCs), neutrophils, and natural killer (NK) cells, forming granuloma. This granuloma contains the Mtb infection and protects from the immune action of lymphocytes.^2^ After the infection is established, Mtb relocates from the lungs to the lymphatics, spreads to other organs and causes extra-pulmonary tuberculosis (EPTB).^1^

In addition to the involvement of these immune cells, emerging evidence demonstrates that Mtb adheres, invades and replicates within the alveolar epithelial cells (AECs).^3–6^ A higher Mtb replication rate is observed in AECs, and such infection leads to host cell death and impairs the epithelial barrier, potentially contributing to the spread from the lungs.^7^ AECs also regulate innate immunity by producing pro-inflammatory cytokines, growth factors and chemokines that interact with immunocytes to activate adaptive immunity.^8,9^ So, Mtb exploits AECs as a “safe haven” to expand and further disseminate during primary infection.^10^ However, studies elucidating the interaction between AECs and Mtb are limited.^10,11^ Understanding the effect of Mtb infection on AECs might enhance our insights into the pathogenesis and immunological mechanisms underlying TB.

Mtb changes its metabolism from active replication to a dormant, non-replicating state to adapt to the host cellular niches. Upon infection, Mtb accumulates mycolic and fatty acids on its cell wall to survive the hostile phagosome environment.^12^ For its nutritional demand, mycobacteria alters the host’s lipid and cholesterol metabolism.^13^ Use of the heavy isotope labelled nutrients (^13^C glucose, glutamine, etc.) helps to identify and understand the perturbed metabolic pathways.^14^

This study aimed to investigate the growth and metabolic response of AECs infected with laboratory and clinical Mtb isolates using 13C-glucose feeding and metabolomics. Specifically, the mass isotopomer distribution of proteinogenic amino acids derived from ^13^C glucose-fed AECs is used as readouts of kinetic metabolic response to decipher the host-pathogen interaction at biochemical levels.

## Results

### Laboratory strains of *Mycobacterium tuberculosis* established infection in the Human alveolar epithelial cell line, A549

A549, a human type 2 alveolar epithelial cell line, and its interaction with Mtb were investigated. A549 cells cultured in Advance DMEM/F12 flex media supplemented with glucose (U-^12^C_6_ or U-^13^C_6_) were infected with virulent (H37Rv) and avirulent (H37Ra) laboratory Mtb strains at MOI of 1:10 (cell: bacteria). Uninfected and heat-killed H37Rv-infected A549 cells were used as controls. Mtb-infected and control cells were harvested at specified time points (i.e., 0, 12, 24, 36, 48, and 60 hours post-infection, h.p.i.) for monitoring different parameters (Figure 1A). Mtb infection impacts host cell viability, affecting disease progression and outcome. Trypan blue assay showed similar viability between uninfected and heat-killed Mtb-infected A549 cells. Virulent Mtb H37Rv infection led to lower A549 cell viability than the H37Ra strain (Figure 1B). Axenic H37Rv and H37Ra cultures showed similar growth patterns in 7H9 broth media (Figure 1C). As intracellular Mtb duplication leads to cell death and release to the culture media which are lost during washing steps, the CFU data was normalised to the number of viable cells (Figure 1D). Both H37Ra and H37Rv showed intracellular survival over 48 h.p.i. in A549 cells, and the CFU was significantly higher upon H37Rv infection than H37Ra at 36 and 48 h.p.i.. Confocal microscopy results showed that the bacteria were distributed majorly around the nucleus close to the mitochondria in a cluster or string-like formations (Figure 1E). H37Rv Mtb strains exhibited higher infection efficiency on A549 cells than avirulent H37Ra. The percentage of A549 cells infected with the H37Ra decreased from 37.5 to 20.5, whereas it increased from 58.4 to 78.6 during 12 to 48 h.p.i. upon H37Rv-infection, corroborating the CFU results. As only two biological replicates were used, statistical significance calculation was not calculated (Figure 1F).

**Figure 1.**
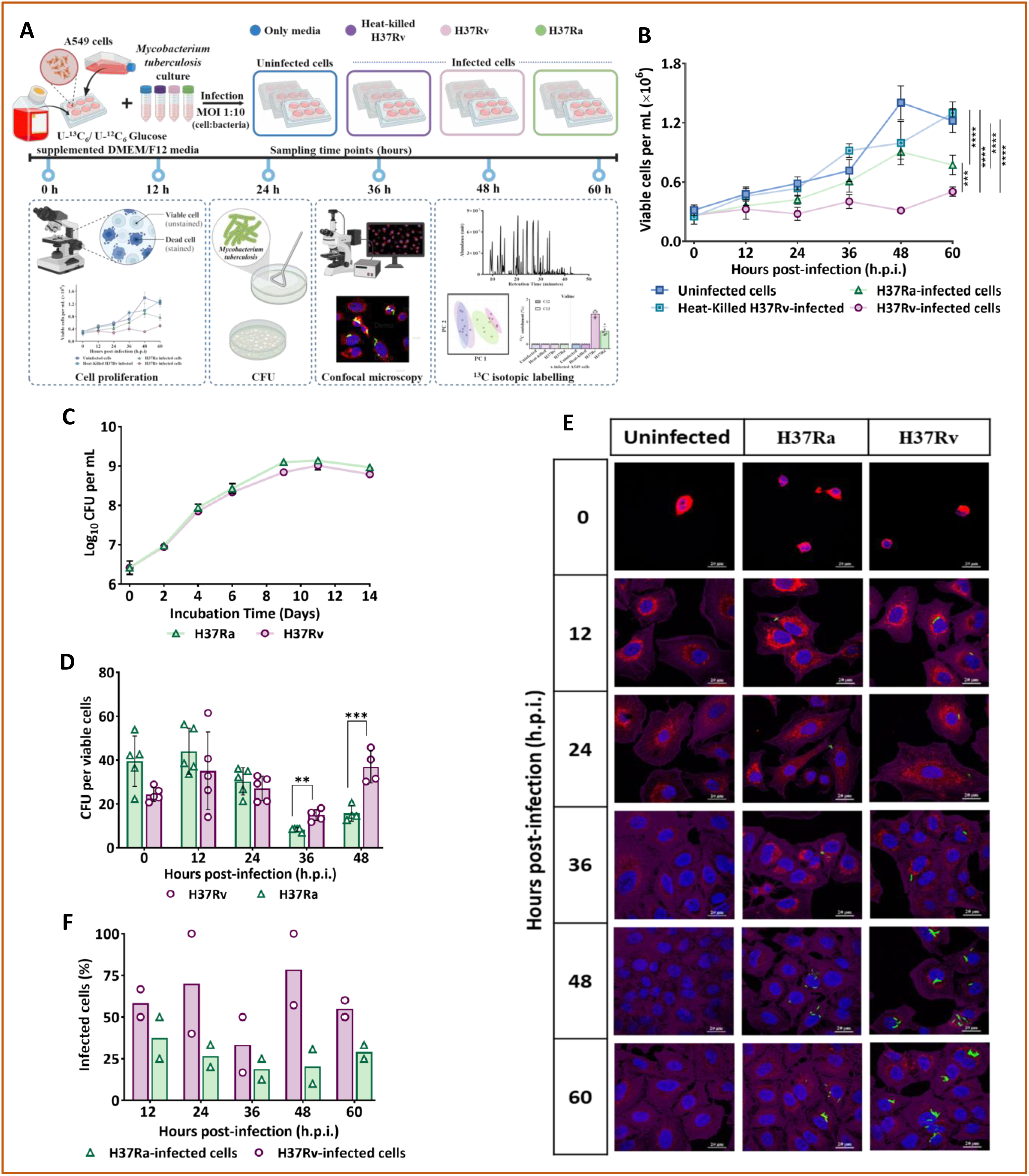
Laboratory *Mycobacterium tuberculosis* strains infect the human alveolar epithelial cell line, A549. (A) Schematic representation of the experimental design of the study. (B) Cell viability of virulent, avirulent and heat-killed Mtb-infected A549 cells with uninfected control. *n* = 5. Error bars represent Standard deviation. (C) Growth rate of Mtb laboratory strains in 7H9 broth media expressed as Log_10_ CFU per mL. *n* = 3. Mean values were presented. (D) Intracellular survival and growth of *Mycobacterium tuberculosis* in A549 cells. Data was plotted as total CFU/total number of viable cells. *n* = 5. (E and F) Human alveolar epithelial cell line (A549) stained with MitoTracker red (red) and Alexa Fluor 680 phalloidin (violet) and DAPI (blue) were infected with laboratory Mtb strain H37Rv and H37Ra respectively (GFP-labelled, green) for 8 hours at an MOI of 1:10, fixed and imaged by (E) confocal microscopy from 0 hour post-infection (i.e. after 8 hours of infection) to 60 hours post-infection. Scale bar, 20 µm. (F) Percentage of A549 cells infected with Mtb strains. *n* = 2. Statistics: one-way ANOVA followed by Tukey’s multiple comparisons test (B), unpaired t-test (C), multiple t-tests with Holm-Sidak correction (D). ^∗∗∗∗^: p < 0.0001; ^∗∗∗^: p < 0.0005; ^∗∗^: p < 0.005; ^∗^: p < 0.05 at 95% confidence interval. A non-significant p-value was not

### Overview of identified amino acids among all study samples

The metabolic interaction between host and pathogen significantly influences the outcome of the infection. Several studies revealed alteration in host cell metabolism as a response to Mtb infection.^15–17^ However, during the infection of AECs with different Mtb strains, the variations in the metabolic response, if any, are not studied in detail. The kinetic isotopic labelling of proteinogenic amino acids can retro-biosynthetically map the activities of central metabolic pathways and shed light on the *de novo* protein biosynthesis. Proteinogenic amino acids are stable and present in abundance inside the host cells, so we aimed to monitor the ^13^C enrichment for a better understanding of the flow of the central metabolic precursors during the infection process. A549 cells were infected with H37Rv, H37Ra and controls (uninfected and heat-killed) and cultured in U-^13^C_6_ glucose to extract proteins at different time points (0, 12, 24, 36, 48 and 60 h.p.i.). Harvested proteins were acid hydrolysed, and analysed using GC-MS to elucidate the average ^13^C-label in the proteinogenic amino acids from the U-^13^C_6_ glucose (Figure 2A). In mammalian cells, most proteinogenic amino acids, except the essential amino acids, are synthesized from the central carbon metabolism, i.e. glycolysis, the pentose phosphate pathway (PPP), and the tricarboxylic acid (TCA) cycle (Figure 2B). A total of 16 proteinogenic amino acids (Alanine, Glycine, Valine, Leucine, Isoleucine, Proline, Methionine, Aspartate/Asparagine, Glutamate/Glutamine, Serine, Cysteine, Histidine, Tyrosine, Lysine, Threonine and Phenylalanine) were identified. Due to chemical decomposition during acid hydrolysis, tryptophan and arginine were undetected. Amino acid-derived mass isotopomer fragments of the ^13^C labelled cells were compared with those derived from the parallel unlabelled (^12^C control) samples. None of the fragments for leucine and isoleucine were found to be valid, so excluded from the comparative study (Table S2). As amino acid pairs like glutamate and glutamine, aspartate and asparagine, could not be distinguished, they were combined as glutamate/n and aspartate/n, respectively.

**Figure 2.**
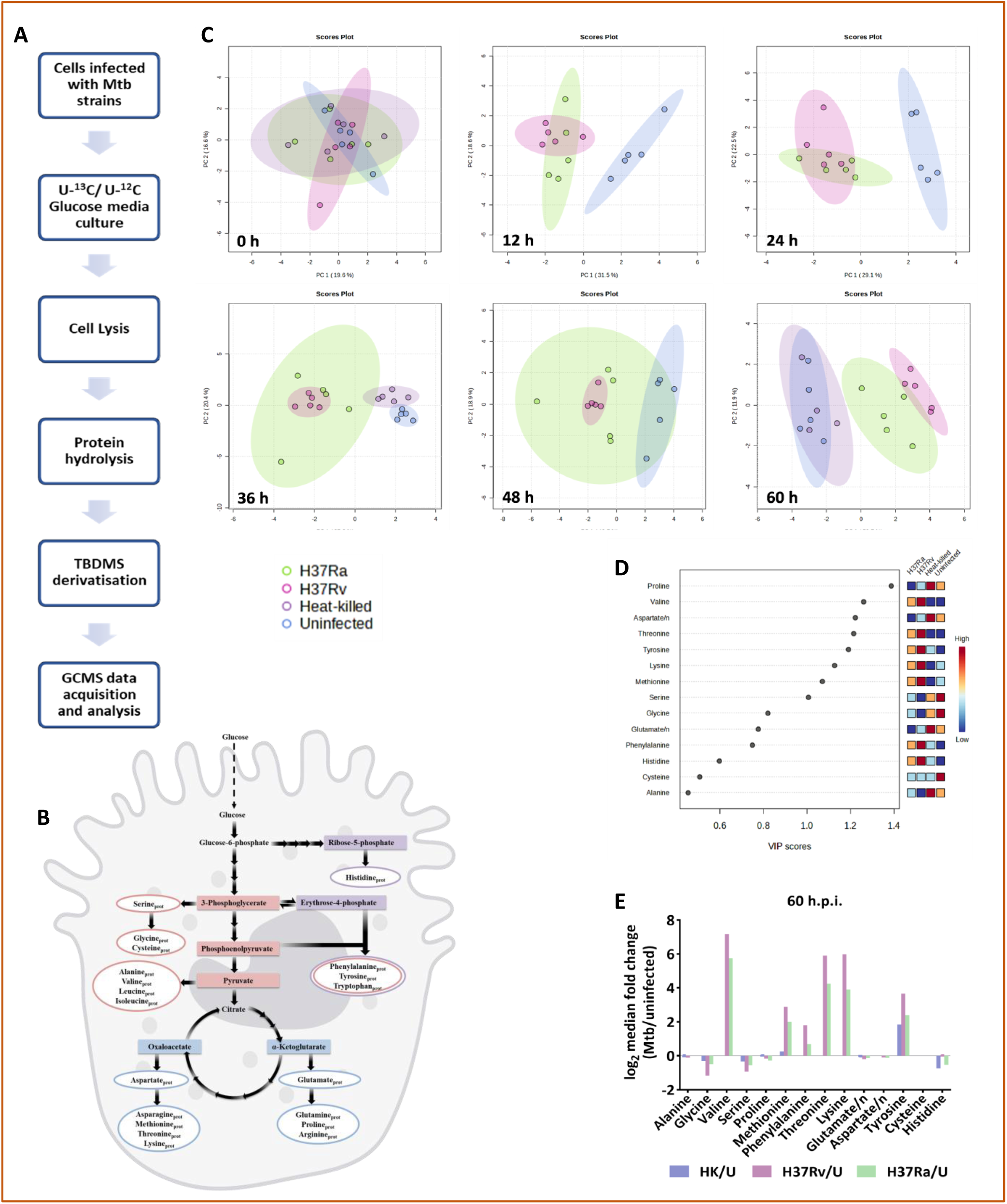
Overview of identified amino acids among all study samples. (A) Schematic of the 13C isotope labelling workflow employed/used for GC-MS-based metabolomics. (B) Schematic illustration of the proteinogenic amino acid biosynthesis pathway. (C) Kinetic Principal component analysis (PCA) plots based on 13C incorporation in the proteinogenic amino acids. (D) Variable Importance for Projection (VIP) scores obtained from the PLS-DA model at 60 h.p.i. (E) Log_2_-transformed proteinogenic amino acid fold change in Mtb-infected versus uninfected A549 cells at 60 h.p.i. The lines represent

The Principal Component Analysis (PCA) of ^13^C incorporation in the proteinogenic amino acid at 60 h.p.i. demonstrated a high similarity between the metabolite profiles of uninfected (U) and heat-killed (HK) H37Rv-infected A549 cells. However, the H37Ra- and H37Rv-infected A549 cells formed distinct clusters (Figure 2C). Eight proteinogenic amino acids (Proline, Valine, Aspartate/n, Threonine, Tyrosine, Methionine, Lysine and Serine) showed a Variable Importance in Projection (VIP) scores >1.0, from the Partial Least Squares Discrimination Analysis (PLS-DA), and selected as important (Figure 2D). Median log_2_ fold change of Mtb-infected compared to uninfected A549 cells at 60 h.p.i. demonstrated changes in the ^13^C enrichment of proteinogenic amino acids upon infection (Figure 2E). The changes in proteinogenic amino acid abundance among groups were significantly different at the later time, i.e. 60 h.p.i.

### Monitoring average ^13^C enrichment kinetics in the proteinogenic amino acids derived from Mtb-infected and uninfected A549 cells provided insights into the central carbon metabolism

Most proteinogenic amino acids showed isotopic labelling post-24 h.p.i.. Therefore, we focussed on monitoring the ^13^C enrichment (%) till 60 h.p.i. to capture maximum differences (Figure 3). Proteinogenic amino acids (alanine, serine, and glycine) synthesized from the precursor of glycolysis intermediates showed varied levels of labelling between Mtb-infected and uninfected A549 cells. The average ^13^C incorporation of Alanine (m/z 260) derived from H37Rv infected cells was significantly less than the uninfected controls as well as those from the heat-killed ones. The average ^13^C incorporation in Serine (m/z 390) was lower in H37Rv- and H37Ra-infected cells compared to HK-H37Rv infected and uninfected control A549 cells. Glycine is synthesized from serine as one of the precursors,^18^ and the proteinogenic glycine (m/z 288) depicted a similar labelling pattern to serine. Uninfected A549 cells showed maximum ^13^C enrichment, followed by HK-H37Rv-infected cells. H37Rv-infected cells exhibited lower ^13^C levels in glycine than H37Ra-infected cells.

**Figure 3.**
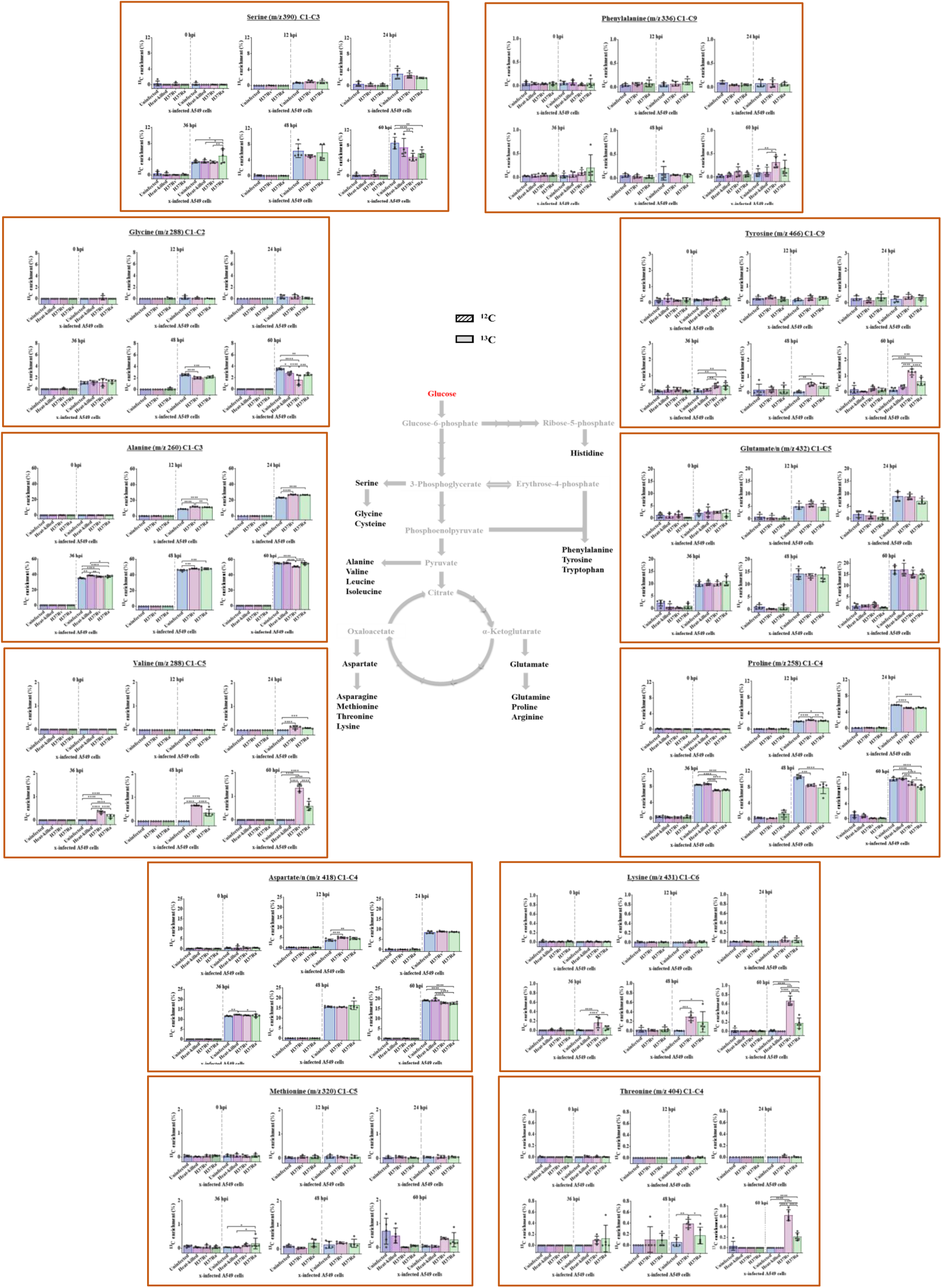
Kinetic ^13^C enrichment (%) into proteinogenic amino acids derived from central carbon metabolic pathway (i.e. Glycolysis, Average ^13^C-isotopic labelling was compared in proteinogenic amino acids harvested from the Mtb-infected (Laboratory strains H37Rv and H37Ra, Heat-killed H37Rv) and uninfected A549 cells. Labelling was compared with parallel unlabelled (^12^C control) samples. The numbers in the bracket beside the amino acid name represent the respective amino acid fragment. Error bars represent the standard deviation of biological replicates (*n* = 5). Statistics: two-way ANOVA followed by Tukey’s multiple comparisons. ^∗∗∗∗^: p < 0.0001; ^∗∗∗^: p < 0.0005; ^∗∗^: p < 0.005; ^∗^: p < 0.05 at 95% confidence interval. A non-significant p-value was not indicated in the plots.

The average 13C levels in the proteinogenic-derived valine, phenylalanine and tyrosine were less than the expected natural abundance of ∼1.13% average ^13^C in the majority of the cases. The contribution of glucose to *de novo* proteinogenic valine (m/z 288) was significantly more in both H37Rv- (1.35±0.16%) and H37Ra (0.59±0.19%) infected cells in comparison to the HK-H37Rv-infected cells and uninfected controls. Phenylalanine and tyrosine are synthesized from the precursors generated from glycolysis and the pentose phosphate pathway (PPP). The Mtb-infected cells acquired the ^13^C levels in Phenylalanine (H37Rv: 0.32±0.12%; H37Ra: 0.19±0.17%) and Tyrosine (H37Rv: 1.24±0.18%; H37Ra: 0.66±0.25%), which was less than the HK-H37Rv-infected cells and uninfected controls.

TCA cycle intermediates contribute to glutamate/n (m/z 432), proline (m/z 258), aspartate/n (m/z 418), methionine (m/z 320), threonine (m/z 404) and lysine (m/z 431). The ^13^C label incorporation of these proteinogenic amino acids provided insights into the contribution of the TCA cycle towards *de novo* protein synthesis (Figure 3). ^13^C incorporated glutamate/n, and proline was observed in all study groups, and they followed a similar pattern. ^13^C incorporation into glutamate/n was observed in all four groups, i.e., Mtb-infected and uninfected control. Live Mtb-infected A549 cells showed slightly lower ^13^C labels than the controls (HK-infected and uninfected A549 cells); however, no significant differences among the groups were observed (uninfected:17.03±1.82%; HK-H37Rv:17.16±2.83%; H37Rv:15.14±1.75%; and H37Ra infected cells:14.84±1.60%). Proteinogenic proline from Mtb-infected A549 cells had significantly lower average ^13^C enrichment (H37Rv-infected cells: 9.38±0.47%; and H37Ra-infected cells: 8.43±0.50%) compared to HK-H37Rv-infected (10.68±0.21%) and uninfected (10.56±0.46%) control A549 cells. Proteinogenic aspartate/n had higher fractions of labelled carbon in uninfected (19.15±0.25%) and HK-H37Rv-infected (19.53±0.72%) A549 cells compared to Mtb-infected A549 cells (H37Rv: 17.95±0.37%; H37Ra: 17.71±0.54%). Interestingly, HK-H37Rv infected and uninfected control A549 cells demonstrated no detectable ^13^C labelling in any of the three aspartate-derived amino acids i.e. methionine, threonine and lysine. These proteinogenic amino acids in Mtb-infected A549 cells showed marginally higher average ^13^C incorporations in comparison to uninfected control cells. Virulent laboratory Mtb strain infected A549 cells showed the average 13C levels in threonine (H37Rv: 0.62±0.10%; and H37Ra: 0.21±0.07%) and lysine (H37Rv: 0.67±0.11%; and H37Ra: 0.18±0.11%). Proteinogenic methionine of H37Rv-infected A549 cells showed similar labelling (H37Rv: 0.38±0.04%; H37Ra: 0.33±0.30%) (Figure 3).

The average ^13^C labelling data suggested that glucose contributed to the *de novo* protein synthesis of A549 cells by continued metabolic activities post-infection. These data also suggested that after 48 h.p.i. (and at 60 h.p.i), variations in ^13^C levels of several *de novo* proteinogenic amino acids derived mainly from the pathway precursors emanating from glycolysis (Alanine, Serine and Glycine) and TCA cycle (Glutamate/n, Aspartate and proline) were captured. Analysis of the mass isotopomer distributions (MIDs) of all the amino acids is further warranted and is presented in the sections below.

### Comparative Mass isotopomer distribution in proteinogenic amino acids of A549 cells upon infection with laboratory *Mycobacterium tuberculosis* strains

Next, we evaluated and compared the MIDs of proteinogenic amino acids derived from the Mtb-infected A549 cells and controls (60 h.p.i.) subjected to [U-^13^C_6_] glucose (Figure 4). The MIDs of the amino acid fragment ions were corrected for the contributions of natural ^13^C isotope abundances. Mtb-infected A549 cells indicated lesser ^13^C labelling in the serine and glycine, as reflected by the higher fractional M+0 (unlabelled pool) compared to the uninfected control cells. The M+3 abundance in alanine was significantly lower in H37Rv-infected A549 cells than in other groups (H37Ra or HK- or uninfected). This is also reflected by a higher abundance of unlabelled M+0 in Alanine of H37Rv-infected cells. In the infected cells, these observations of higher unlabelled M+0 (and lower abundances of any other isotopomers) in glycine and alanine can be attributed to either slower rate of *de novo* protein biosynthesis or higher contributions of pre-existing or exogenous metabolic intermediates that contribute to these soluble amino acid pools via the glycolytic pathways. Contrary to the patterns in alanine, ^13^C incorporation in valine was higher in both H37Rv and H37Ra Mtb-infected cells than in uninfected or heat-killed controls. This is reflected by the lower levels of unlabelled M+0 and higher levels of M+4 and M+5. No significant differences in the levels of glutamate/n mass isotopomers was observed among the Mtb-infected or uninfected controls. However, H37Rv Mtb infection exhibited higher unlabelled M+0 and lower ^13^C levels of other MIDs in aspartate (M+2) and proline (M+2 and M+3) compared to the controls (uninfected, HK-infected cells). These variations in the ^13^C labelling of aspartate and proline indicate possible differences in the TCA cycle activities in the Mtb-infected cells. Mtb infection of A549 cells induced significant changes in the MIDs of tyrosine (, M+4, M+5 and M+6) but not of phenylalanine (M+5), which also points to possible variation in the pentose phosphate pathway activities in Mtb-infected cells. Interestingly, in the Mtb-infected cells, the lower levels of M+0 of threonine, methionine and lysine fragments point to the ^13^C label incorporation, as can be confirmed from the higher levels of other mass isotopomers due to ^13^C incorporation (threonine M+3 and M+4; methionine M+3 and lysine M+3, M+4, and M+5). These observations suggest the active metabolism of intracellular Mtb to biosynthesise its amino acids by harnessing the metabolic intermediates formed by the infected cells using substrates available in exogenous growth media.

**Figure 4.**
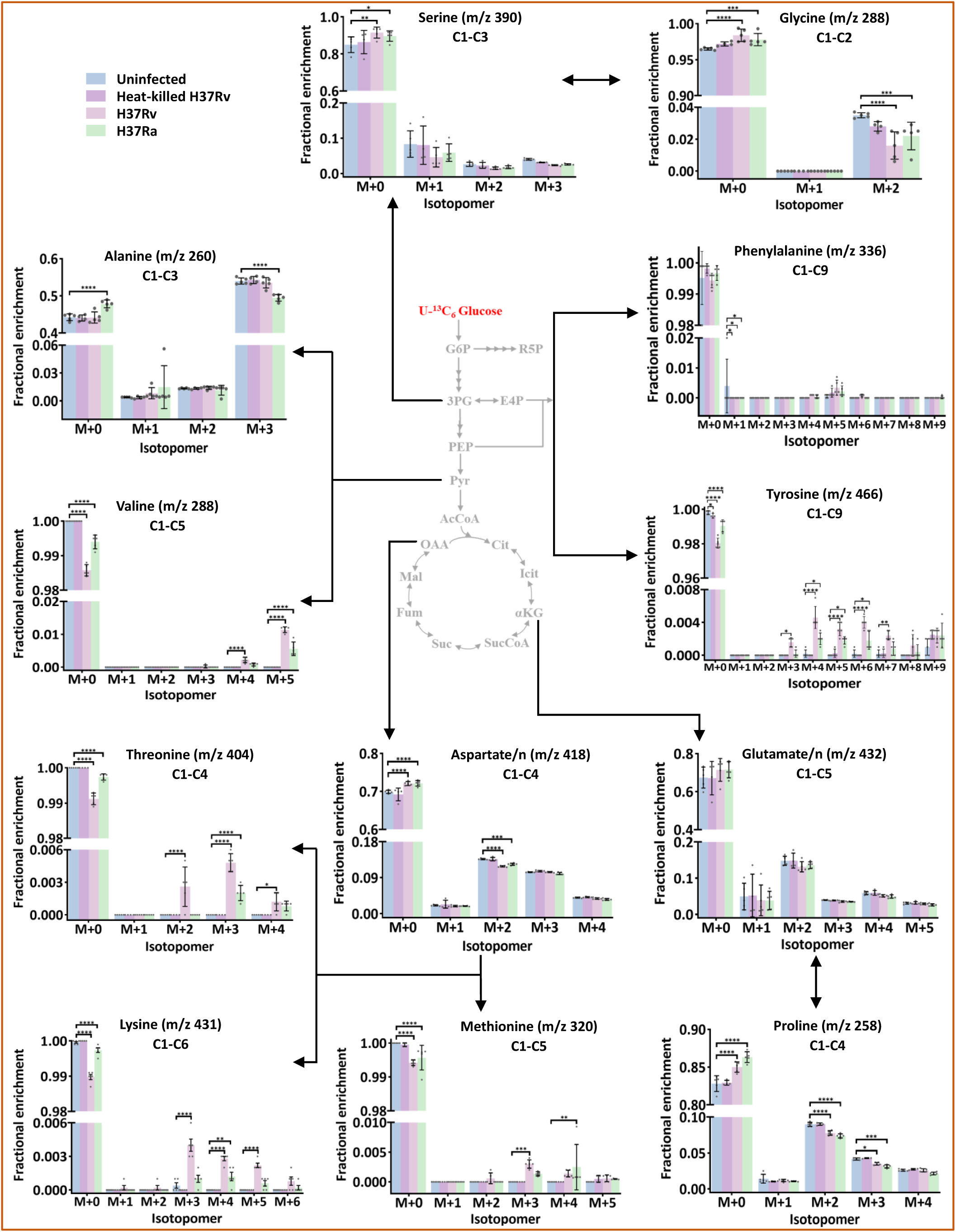
Comparative ^13^C mass isotopomer distributions (MIDs) in proteinogenic amino acids of A549 cells infected with Mtb at 60 hours post-infection. M+0, M+1, M+2, M+3, M+4, M+5, M+6, M+7, M+8, and M+9 are the mass isotopomers of the amino acids. Error bars represent the standard deviation of biological replicates (*n* = 5). Statistics: Two-way ANOVA with Dunnett’s multiple comparisons test. Multiplicity adjusted p-value: ^∗∗∗∗^: p < 0.0001; ^∗∗∗^: p < 0.0005; ^∗∗^: p < 0.005; ^∗^: p < 0.05 at 95% confidence interval. A non-significant p-value was not indicated in the plots.

### Clinical *Mycobacterium tuberculosis* (Mtb) isolates successfully infect the Human alveolar epithelial cells

The rising emergence of multi- and extensively drug-resistant Mtb strains (MDR and XDR) presents significant challenges to TB management. Understanding metabolic pathways specific to drug-resistant strains might provide effective novel treatment strategies.^19^ Therefore, we focused next on comparing the interaction of A549 cells with well-characterized drug-sensitive (S4 and S5) and drug-resistant (MDR: S6 and XDR: S11) clinical Mtb isolates infected with an MOI of 1:10. A549 cells were cultured in Advance DMEM/F12 flex media supplemented with glucose (100% U-^12^C_6_ /U-^13^C_6_) and the proteinogenic amino acids were harvested at specified time points (i.e., 0, 24 and 48 h.p.i.) were analyzed (Figure 5A). Mtb H37Rv, clinical S4 and S5 isolates were sensitive to Isoniazid (INH), Rifampicin (RIF), Streptomycin (STM), Moxifloxacin (MOX), and Kanamycin (KAN). The clinical Mtb isolate, S6 was resistant to INH, RIF, and STM, and S11 was resistant to STM, INH, and MOX (Figure 5B and Table S1). These clinical Mtb isolates showed similar growth kinetics up to 4 weeks in 7H9 broth media (Figure 5C). However, strain-specific differences in intracellular survival and replication, like CFU per viable cells, were observed. The S4, S5, and S11 clinical Mtb isolates exhibited low mycobacterial burden in the early time points, whereas at 48 h.p.i., intracellular bacilli increased, as observed from the CFU data. S4 data at 48 h.p.i. could not be reported due to contamination. H37Rv and S6 Mtb strains showed a six- and eighteen-fold increase in intracellular bacillary load at 48 h.p.i. compared to early time point (0 h.p.i.). Compared to uninfected A549 cells, Mtb infection decreased the viability of these cells over time, and H37Rv Mtb strain infected cells showed the lowest viability compared to the clinical isolates. Interestingly, A549 cells infected with clinical MDR or XDR Mtb isolates exhibited higher viability and a higher number of Mtb accumulation than those infected with drug-sensitive clinical isolates (Figures 5E and 5F). At the selected MOI of 1:10, most (80-100%) of the A549 cells were infected. Hence, the observed perturbed phenotypes may largely be attributed to the drug resistance patterns (Figure 5G).

**Figure 5.**
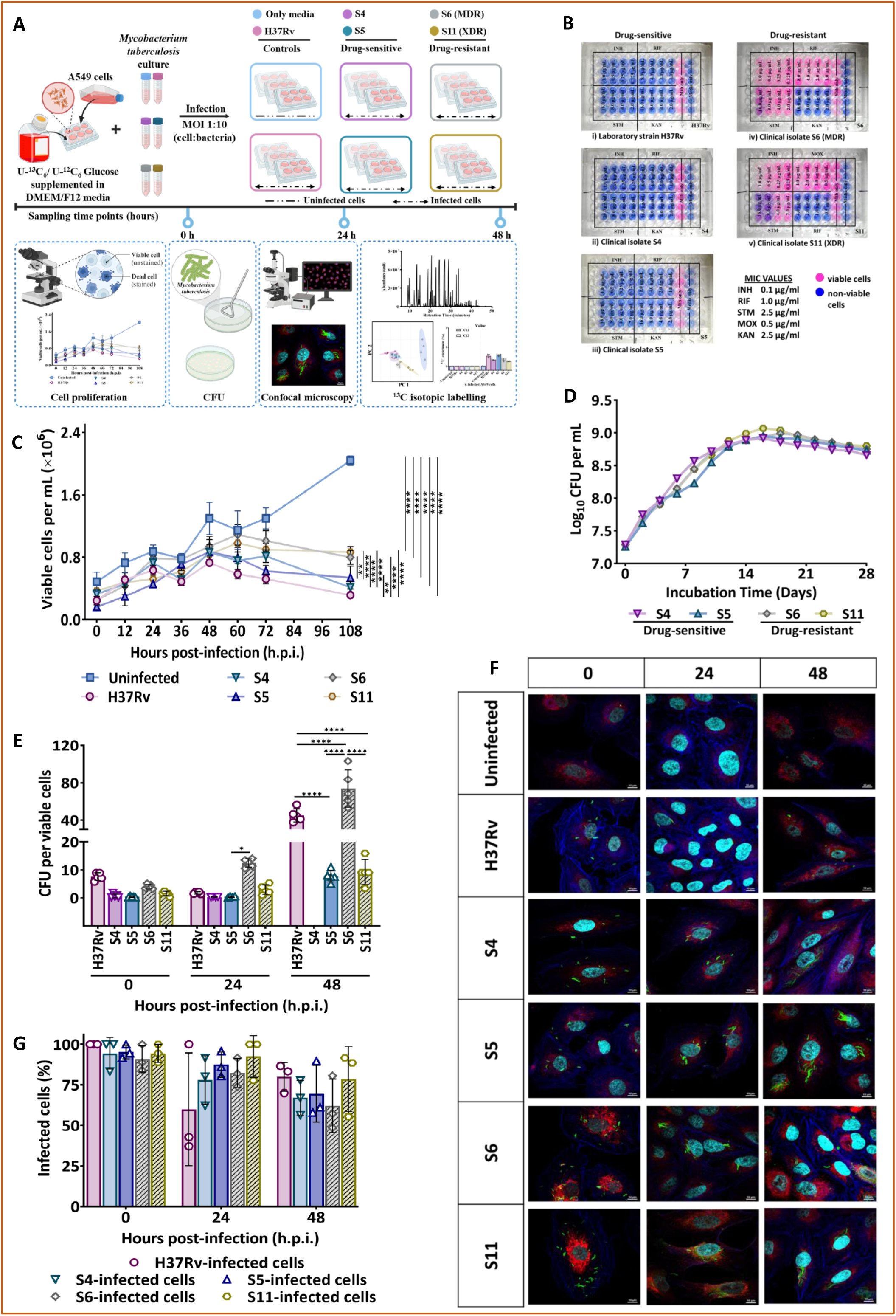
Clinical *Mycobacterium tuberculosis* isolates successfully establish infection in the Human alveolar epithelial cell line, A549. (A) Schematic representation of the experimental protocol. (B) Alamar Blue antibiotic susceptibility assay of laboratory strain H37Rv and clinical isolates S4, S5, S6, and S11. TB drugs: Isoniazid (INH), Rifampicin (RIF), Streptomycin (STM), Moxifloxacin (MOX), and Kanamycin (KAN). (C) Cell viability of Mtb-infected A549 cells with uninfected control. *n* = 5. Error bars represent Standard deviation. (D) Intracellular survival and growth of *Mycobacterium tuberculosis* in A549 cells. Data is plotted as total CFU/total number of viable cells. S4 data was not included in determining statistical significance as 48-hour data could not be recorded. *n* = 5. (E) Growth rate of (Mtb) clinical isolates in 7H9 broth media. *n* = 3. Mean values are presented. (F and G) Human alveolar epithelial cell line (A549) stained with MitoTracker red (red) and Alexa Fluor 680 phalloidin (false colour blue) and DAPI (false colour cyan) were infected with Mtb reference laboratory strain H37Rv and clinical isolates (PKH67-stained, green) for 8 hours at an MOI of 1:10, fixed and imaged by (F) confocal microscopy from 0 hour post-infection (i.e. after 8 hours infection) to 48 hours post-infection. Scale bar, 10 µm. (G) Percentage of A549 cells infected with Mtb strains. *n* = 3. Statistics: one-way ANOVA followed by Tukey’s multiple comparisons test (C and D); two-way ANOVA followed by Tukey’s multiple comparisons test (E - G). ^∗∗∗∗^: p < 0.0001; ^∗∗∗^: p < 0.0005; ^∗∗^: p < 0.005; ^∗^: p < 0.05 at 95% confidence interval. A non-significant p-value was not indicated in the plots.

### Qualitative insights into the variations in the ^13^C enriched proteinogenic amino acids from alveolar epithelial cells (A549) upon infection with laboratory strain H37Rv and clinical isolates of Mtb

The Principal component analysis (PCA) of the ^13^C enriched fragments ions of proteinogenic amino acids among the study groups demonstrated separate clusters of the uninfected and the Mtb-infected A549 cells prominently after 48 h.p.i. A549 cells infected with drug-sensitive clinical isolates exhibited some overlap while forming a distinct group from the H37Rv-infected cells. Notably, cells infected with drug-resistant clinical isolates clustered distinctly away from other infected groups, highlighting the potential impact of the pathogen’s drug-resistance phenotype on the perturbed host cellular response (Figure 6A). Based on VIP scores >0.5, calculated from the PLS-DA model, eleven amino acids seemed to contribute to the variations at 48 h.p.i (Figure 6B). Additionally, the median log_2_ fold change of mean ^13^C enrichment in amino acid fragments of Mtb-infected versus uninfected A549 cells at 48 h.p.i. exhibited variations that merit further investigation (Figure 6C).

**Figure 6.**
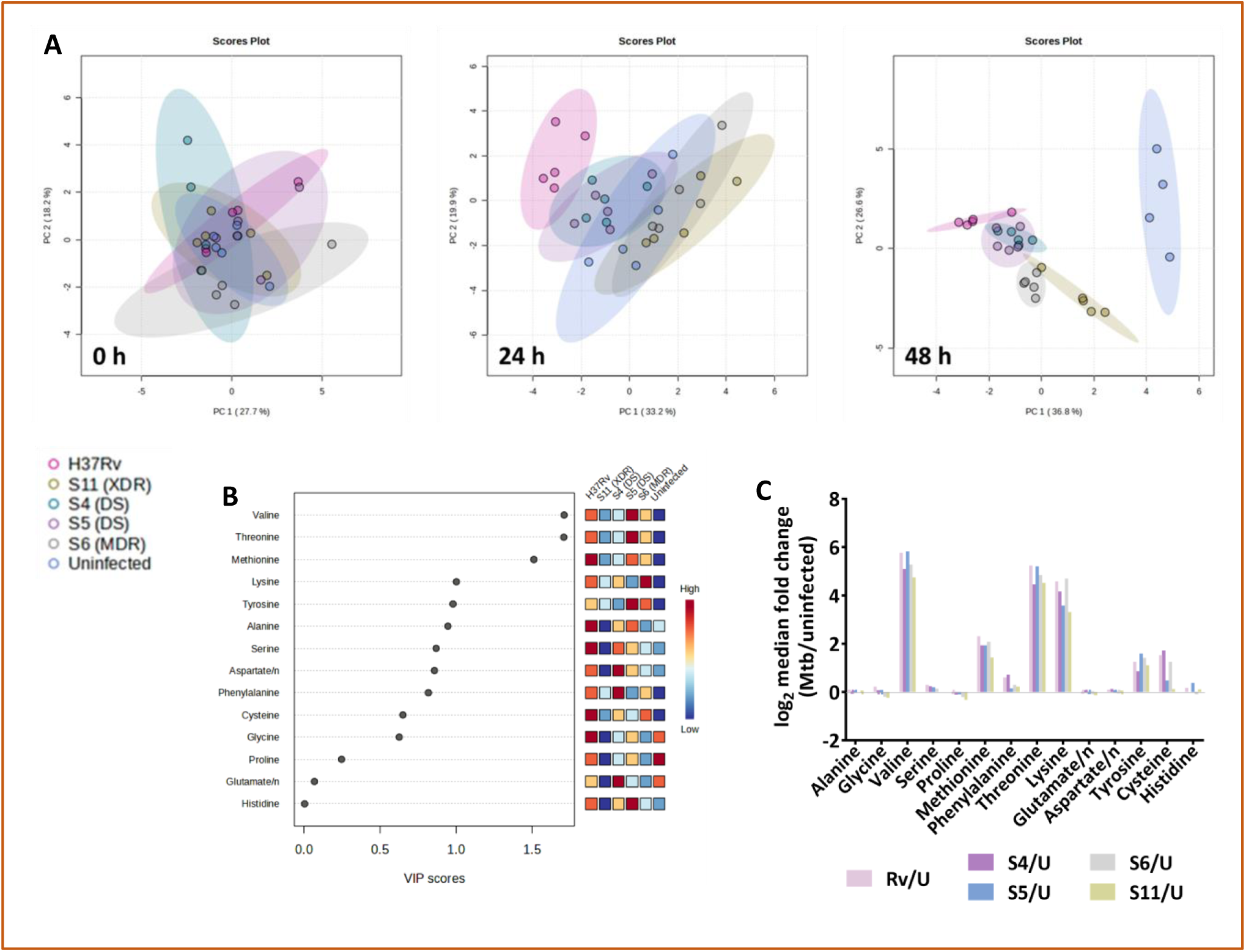
Metabolic Profiling of identified amino acids from alveolar epithelial cells upon infection with laboratory strain H37Rv and clinical isolates of Mycobacterium tuberculosis (Mtb) (A) Principal component analysis (PCA) plots based on ^13^C enrichment in proteinogenic amino acids across time points. *n* = 5 per group. (B) Variable Importance for Projection (VIP) scores at 48 hours post-infection (h.p.i.). (C) Amino acids sorted by average median fold change (C) at 48 h.p.i. The lines represent the median log_2_ fold change relative to baseline. Bars represent group medians.

### ^13^C enrichment in the proteinogenic amino acids derived from alveolar epithelial cells (A549) infected with laboratory Mtb H37Rv strain and clinical isolates provides insights into the central metabolism and the dependence of exogenous amino acids

Proteinogenic amino acids extracted from the U-^13^C_6_ glucose-supplemented A549 cells infected with laboratory Mtb H37Rv strain and clinical isolates at 0, 24 and 48 h.p.i. were monitored to ascertain the average ^13^C-label.

Among all the proteinogenic amino acids, average ^13^C labelling of alanine (m/z 260) at 48 h.p.i., was observed to be the highest and exhibited enrichment in drug-sensitive Mtb-infected cells (incuding H37Rv) among the uninfected and infected cells (uninfected: 43.88±0.57%; H37Rv: 47.16±0.81%; S4: 44.49±0.36%; S5: 46.04±0.73%; S6: 43.80±0.40%; S11: 42.70±0.76%). Average ^13^C labelling of serine was observed in all groups (uninfected: 8.83±0.89%; H37Rv: 10.36±0.29%; S4: 9.70±0.38%; S5: 9.67±0.20%; S6: 9.20±0.28%; and S11: 8.45±0.68%). Proteinogenic glycine (m/z 288) displayed a similar labelling pattern to serine (uninfected: 3.81+0.10%; H37Rv: 4.34±0.14%; S4: 3.58±0.40%; S5: 3.76±0.16%; S6: 3.38±0.21%; and S11: 3.21±0.10%). A549 cells infected with Mtb clinical isolates (S4, S5, S6 and S11) exhibited less ^13^C enrichment than those infected with the laboratory strain H37Rv. The cells infected with drug-resistant clinical isolates (S6 and S11) showed the least enrichment among all the infected groups. The ^13^C enrichments and the variations observed in the proteinogenic alanine, serine and glycine can be attributed to the changes in the activities of glycolytic pathways and other metabolic reactions in a strain-specific manner.

In uninfected cells, the proteinogenic valine (m/z 288) and tyrosine (m/z 466) showed no detectable ^13^C enrichment (i.e. less than natural abundance of ∼1.13%), probably indicating the contribution of these amino acids from exogenous media. However, when compared with the uninfected controls, the Mtb-infected cells seem to have strain-specific variations albeit at lower levels, which needs a further critical examination of MIDs to infer any metabolic activities.

The incorporation of ^13^C into proteinogenic amino acids from the TCA cycle, i.e. aspartate/n, glutamate/n, methionine, threonine, lysine, and proline, at 48 h.p.i was monitored. It was observed that ^13^C from glucose was incorporated into glutamate/n and aspartate/n in all groups, including uninfected controls. In glutamine/n (m/z 432), the ^13^C enrichment (%) varied to a small extent among the cells (uninfected: 15.31±1.42%; H37Rv: 15.13±0.30%; S4: 15.76±0.48%, S5:14.99±0.38%, S6: 14.07±0.45%; S11: 14.06±0.92%). For aspartate/n (m/z 418), minor differences in ^13^C enrichment (%) was observed (uninfected: 17.45±1.12%; H37Rv: 18.80±0.21%; S4: 19.08±0.27%; S5: 18.61±0.28%; S6: 17.88±0.38%; and S11: 17.33±0.42%). The ^13^C enrichment of proline (m/z 258) derived from the clinical isolates infected A549 cells were significantly lower (S4: 7.74±0.13%, S5: 8.09±0.57%; S6: 7.25±0.39%; S11: 6.76±0.25%) than other uninfected (8.52±0.43%) and H37Rv infected cells (8.50±0.33%). The Mtb-infected A549 cells displayed no considerable ^13^C enrichment in the proteinogenic amino acids derived from aspartate i.e. methionine (m/z 320), threonine (m/z 404), and lysine (m/z 431) and were less than the natural abundance of ∼1.13%. On careful observation of average 13C enrichment, although less than natural abundance, it seems there are significant variations in a strain-specific manner. In the case of methionine, the ^13^C enrichment (%) was different between groups (H37Rv: 0.32±0.07%; S4: 0.26±0.03%; S5: 0.28±0.06%; S6: 0.28±0.04%; S11: 0.18±0.02%). For threonine, the ^13^C enrichment (%) showed Mtb strain specific differences (H37Rv: 0.34±0.11%; S4: 0.24±0.07%; S5: 0.37±0.09%; S6: 0.25±0.08%; and S11: 0.21±0.05%). Varying incorporation of ^13^C label in lysine across groups (H37Rv: 0.21±0.09%; S4: 0.14±0.08%; S5: 0.11±0.08%; S6: 0.27±0.05%; and S11: 0.12±0.07%) was observed. Careful examination of the MIDs is further warranted to make any reasonable conclusions.

A549 cells infected with drug-resistant Mtb clinical isolates showed a minor decrease in the ^13^C label than other infected groups. However, no significant differences were observed in the labelling patterns of A549 proteinogenic amino acid infected with the laboratory H37Rv strain and the drug-sensitive isolates S4 and S5 (Figure 7 and Table S3). These findings point towards drug resistance-pattern-specific selective metabolic adaptations observed in the Mtb-infected type 2 alveolar epithelial cells. Low ^13^C enrichment in a few proteinogenic amino acids likely reflects the contribution of exogenous carbon sources from the host or culture media or preexisting unlabelled metabolite pools of cells before incubation with labelled glucose. Also, the variations in the rates of *de novo* protein biosynthesis and the central metabolic fluxes in the uninfected and infected cells can contribute to the differences. Analysis of MIDs can further shed light on the phenotypes.

**Figure 7.**
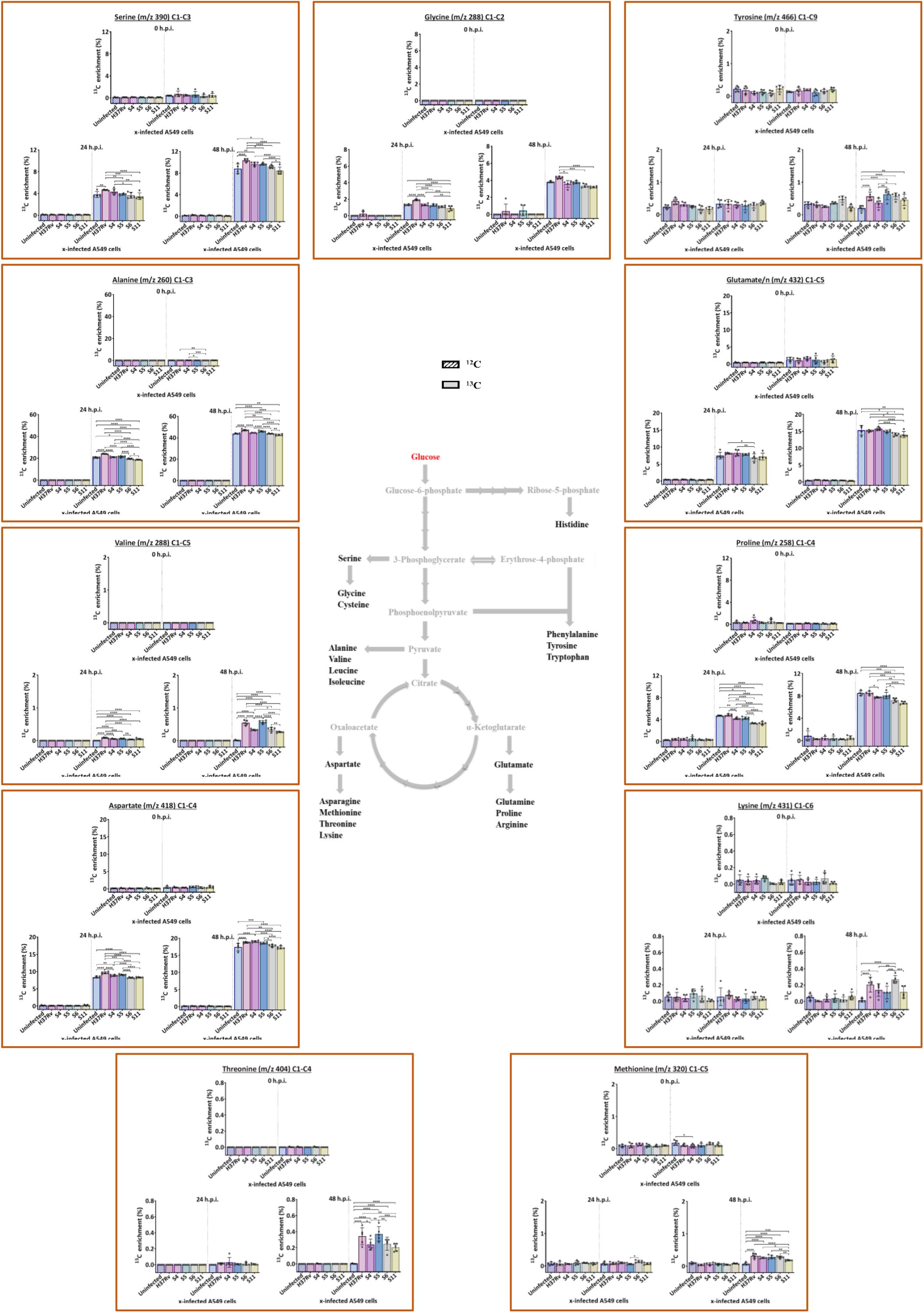
Kinetic ^13^C enrichment (%) of proteinogenic amino acids derived from central carbon metabolic pathway (i.e. Glycolysis, Pentose Phosphate Pathway (PPP) and Tricarboxylic Acid cycle (TCA)) in Mtb-infected and uninfected A549 cells. Average ^13^C-isotopic labelling was compared in proteinogenic amino acids harvested from the Mtb-infected (Laboratory strain H37Rv, drug-sensitive: S4 and S5, and drug-resistant: S6 and S11 clinical isolates) and uninfected A549 cells. Labelling was compared with parallel unlabelled (^12^C control) samples. The numbers written in the bracket beside the amino acid name represent the respective amino acid fragment. Error bars represent the standard deviation of biological replicates (*n* = 5). Statistics: two-way ANOVA followed by Tukey’s multiple comparisons. ^∗∗∗∗^: p < 0.0001; ^∗∗∗^: p < 0.0005; ^∗∗^: p < 0.005; ^∗^: p < 0.05 at 95% confidence interval. A non-significant p-value was not indicated in the plots.

### Comparative mass isotopomer distribution profiles of proteinogenic amino acids derived from A549 cells infected with laboratory Mtb strain H37Rv and the clinical isolates

We compared the MIDs of proteinogenic amino acids obtained from uninfected and Mtb-infected A549 cells at 48 h.p.i. (Figure 8). The labelling pattern of alanine was characterised by a high fractional enrichment in M+3 mass isotopomer in the order of H37Rv > S5 > uninfected∼S4∼S6 > S11 infected cells. The mean proportions of M+1 and M+2 were slightly higher in Mtb-infected A549 cells, but no significant differences were observed among the study groups. In serine, the drug-resistant Mtb clinical isolates (S6 and S11) infection exhibited lower levels of mean ^13^C incorporation compared to H37Rv, and drug-sensitive Mtb infected A549 cells as reflected by the M+0 enrichment. Higher mean enrichment of serine M+3 indicates variable flux from [U-^13^C_6_] glucose via ^13^C_3_-labelled glycolytic intermediates in Mtb-infected cells. Fractional enrichment of the mass isotopomers M+0 and M+2 of glycine in the drug-resistant Mtb clinical isolates (S6 and S11) indicated significantly lower 13C incorporation compared to the uninfected, and drug-sensitive clinical Mtb isolates (S4 and S5) infected cells indicating potential variations in the glycolytic pathway rates among the groups. Infection with drug-resistant Mtb clinical isolates (S6 and S11) seemed to decrease the carbon flow to glycolytic intermediate 3-Phosphoglycerate (3PG)-derived glycine. The fractional enrichment of valine M+0 and M+5 showed higher ^13^C incorporation in Mtb-infected A549 cells compared to the uninfected controls. The abundance of valine M+5 was higher in H37Rv-infected cells, followed by S5-infected cells. XDR S11-infected cells incorporated significantly lower ^13^C among the infected groups. In cells infected with the drug-sensitive clinical isolate S5, the enrichment of M+4 was the highest. However, the relative quantities in all other infected groups were comparable.

**Figure 8.**
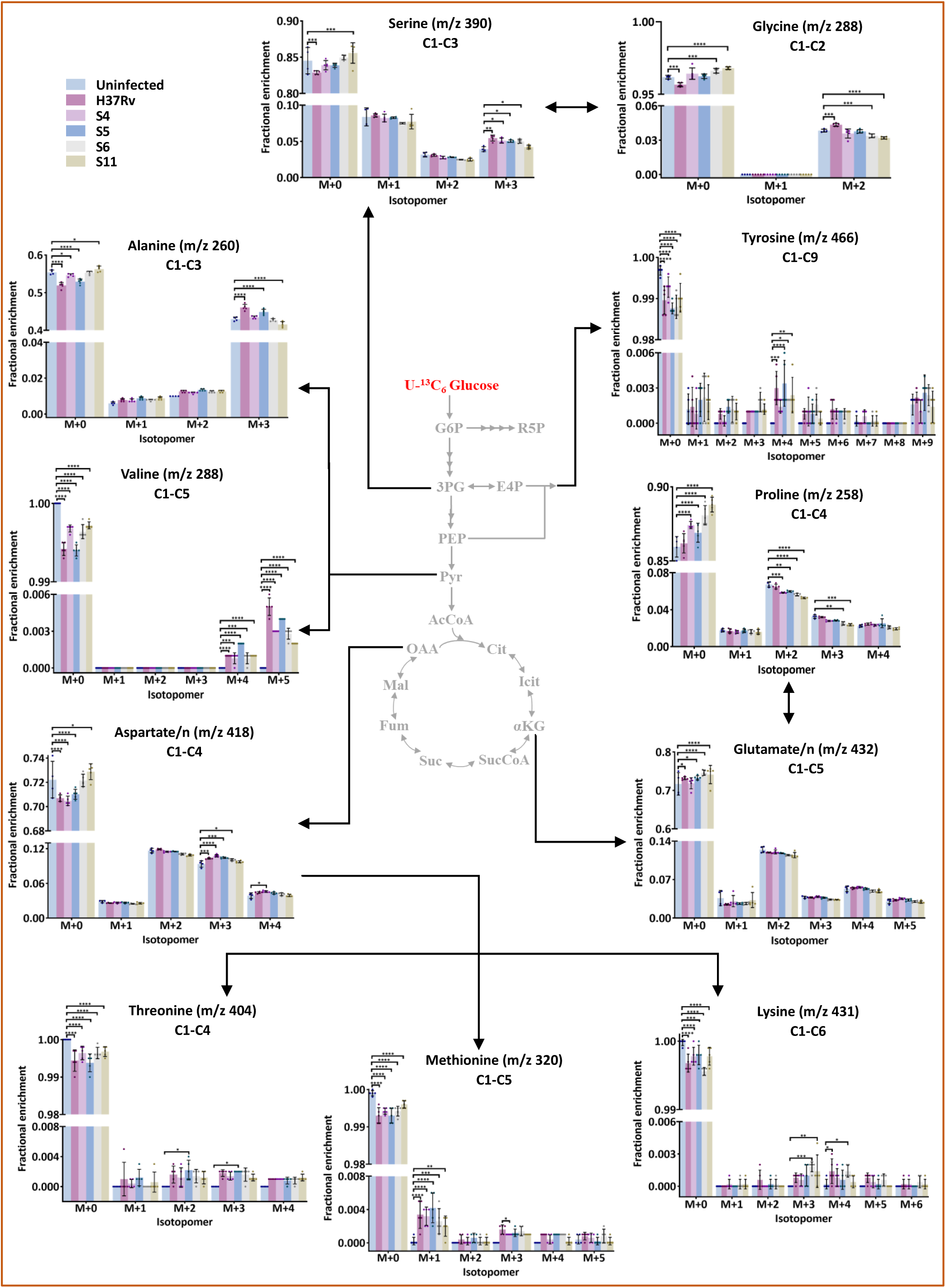
Comparative ^13^C mass isotopomer distributions in proteinogenic amino acids of A549 cells infected with Mtb at 48 hours post-infection. M+0, M+1, M+2, M+3, M+4, M+5, M+6, M+7, M+8, and M+9 are the mass isotopomers of the amino acids. *n* = 5 per group per time-point. Statistics: Two-way ANOVA with Dunnett’s multiple comparisons test. Multiplicity adjusted p-value: ^∗∗∗∗^: p < 0.0001; ^∗∗∗^: p < 0.0005; ^∗∗^:

Tyrosine is synthesised from phenylalanine derived from the Pentose Phosphate Pathway and glycolytic intermediates. Despite low ^13^C incorporation, the M+0 enrichment profile differed between uninfected and Mtb-infected groups indicating variable fluxes through these pathways on infection.

The mass isotopomer M+2 of glutamate/n points to the reduced mean ^13^C enrichment in cells infected with drug-resistant Mtb clinical isolates compared to uninfected control. Fractional enrichment of M+0 of proline showed Mtb infection induced decreased ^13^C incorporation with a significant difference between cells infected with Mtb clinical isolates and uninfected control. Lowest isotopic enrichment was observed in cells infected with drug-resistant clinical isolates compared to other infected groups. Mean fractional enrichment of M+0 aspartate/n points to the lower ^13^C incorporation in drug-resistant Mtb clinical isolates (S6 and S11) compared to other Mtb (H37Rv, S4 and S5) infected cells potentially highlighting strain-specific metabolic adjustments. In methionine, fractional abundances of M+0 and M+1 point to the higher ^13^C incorporation in the infected cells compared to uninfected controls. In both threonine and lysine, fractional abundances of M+0 point to the higher ^13^C incorporation in the infected cells compared to the uninfected controls. In threonine, M+2 and M+3 were higher in all Mtb-infected groups compared to the uninfected controls. Cells infected with drug-resistant Mtb isolates exhibited higher fractional enrichment of M+3 of lysine than drug-sensitive Mtb-infected cells. Strain-specific ^13^C enrichment was observed in the isotopomer M+4 of lysine. This ^13^C enrichment study revealed robustness in the central carbon metabolism of type 2 alveolar epithelial cells due to Mtb infection, irrespective of their origin (laboratory strains or clinical isolates) and depending on their virulence status. Infection-induced ^13^C enrichment in essential proteinogenic amino acids (i.e., valine, threonine, methionine and lysine) was lowest upon infection with drug-resistant clinical Mtb isolates.

## Discussion

In the present study, we aimed to gain insights into the host alveolar epithelial cell type 2 (AEC2) response against Mtb infection of either laboratory-established or clinical strains of varied drug resistance status. We performed a time-dependent comparative evaluation for bacterial growth rate in axenic culture, host cell viability, intracellular bacterial viability, infected cell numbers and proteinogenic amino acids analyses of Mtb-infected A549 cells subjected to ^13^C Glucose-containing media. Heat-inactivated H37Rv served as a control to match the particle size of live Mtb. ^13^C stable-isotope tracing was adopted to quantify the ^13^C enrichment in proteinogenic amino acids obtained from uninfected and Mtb-infected A549 alveolar epithelial cell adenocarcinoma cell line as the *in vitro* infected host model to study the robustness of host-pathogen interactions.

Our observations indicated that live Mtb, particularly the laboratory-adapted virulent H37Rv strain, significantly reduced A549 cell viability, consistent with similar reports on Mtb-infected macrophages and dendritic cells.^20,21^ A549 cells infected with drug-resistant clinical isolates showed greater viability than drug-sensitive ones. These findings signify the differential impact of various Mtb strains on the host cell viability, reiterating that distinct genotypes of bacterial strains, the host’s immune response, and the stress responses induced by the infection may contribute to the observed differences in cell viability.

Mtb H37Ra and H37Rv strains persist and replicate within A549 cells.^5,22,23^ Differential growth of H37Rv within cells can be attributed to the presence of virulence factors like serine protease Rv2569c, which facilitate the translocation of Mtb across the epithelial barrier by disrupting E-cadherin^24^; leading to varying degrees of apoptosis^25^; and the distinct metabolic responses elicited in host cells.^26^ MDR clinical isolate S6 exhibited maximum intracellular growth (eighteen-fold), followed by the H37Rv (six-fold). The S6 clinical Mtb isolate belongs to the Beijing lineage^27^, which may be one of the reasons for its significantly higher bacterial burden relative to the other strains evaluated.^28,29^ Mtb clinical isolates are reported to exhibit rapid proliferation in human macrophages than H37Rv.^30^ These findings suggest that the CFU counts of clinical Mtb isolates in host cells vary significantly depending on the strain’s ability to adapt and survive within the host cellular environment.

The ^13^C incorporation, determined by GC-MS analysis, predominantly occurred in six proteinogenic amino acids in the order alanine > aspartate/n > glutamate/n > proline > serine > glycine. ^13^C enrichment confirmed that heat-killed H37Rv do not engage in significant metabolic perturbation. The observed enrichment primarily represents host metabolites, whereas ^13^C enrichment, if any, in essential amino acids (valine, phenylalanine, threonine, methionine and lysine) indicates their origin from the bacterial fraction, highlighting active metabolic activity and protein synthesis by intracellular Mtb. Our ^13^C enrichment study revealed robustness of the central carbon metabolism of A549 cells during infection with laboratory strains and clinical Mtb isolates, with appreciable perturbations observed in the later time points. Infection with different Mtb virulent strains showed a consistent pattern in the labelling of the proteinogenic amino acids. Despite very low ^13^C labels, cells infected with the H37Rv strain showed increased ^13^C enrichment in the essential proteinogenic amino acids compared to the controls. In contrast, the enrichment was slightly lower in cells infected with drug-resistant Mtb. This reduced isotopic enrichment in the drug-resistant Mtb-infected A549 cells reflects slower metabolic activity. Drug-resistant Mtb tends to enter a dormant state with decreased metabolic processes, allowing it to adapt and survive longer within host cells.^31,32^ Proline significantly increased in A549 cells infected with the drug-sensitive Mtb compared to those infected with MDR and XDR isolates.^33^ The Mtb-infected A549 cells displayed minimal ^13^C labelling in methionine, threonine and lysine in a strain-specific manner. These essential amino acids, produced through the aspartate pathway, synthesize diaminopimelate (DAP) and S-adenosyl methionine (SAM), which are vital for the survival of mycobacteria.^34^ These observations underlined the dependencies of Mtb on the *de novo* synthesised host metabolic precursors for its sustenance.

The metabolic variations among the A549 cells infected with different Mtb strains seem to get more distinct with the infection time. Alterations in the ^13^C isotopomer distribution profiles of alanine, glycine, serine, aspartate/n, glutamate/n, and proline in Mtb-infected A549 cells, suggested that minor variations in the host cell carbon flux through glycolysis and the TCA cycle as a result of Mtb infection. This highlights largely the robustness of the metabolic activities during the intracellular growth of Mtb, possibly a survival strategy to meet its cellular demands. Contrary to findings from a comparative macrophage study, where ^13^C isotopomer distribution in serine showed M+3 predominance in uninfected and M+1 in infected macrophages,^35^ we observed no difference in serine isotopomer distribution between uninfected and infected A549 cells. However, the abundance of M+3 isotopomer increased in the Mtb-infected cells compared to uninfected control cells. These distinct distributions suggest cell-specific metabolic responses to Mtb infection or stimuli. An earlier report on the Mtb-infected macrophages indicated that ^13^C_3_-labelled host pyruvate is not a significant carbon source for intracellular Mtb.^36^ Our work on A549 cells suggested that the M+5 valine isotopomer in Mtb originates from two ^13^C_3_-labelled pyruvate precursors. The tyrosine distribution profile of ^13^C showed notable differences between uninfected and infected groups in a strain-specific pattern.

Serine M+3 isotopomer distribution indicates that it is synthesized from the isotopomer precursors of 3-phosphoglycerate. Another possible source of ^13^C-serine is ^13^C-glycine through the reversible serine hydroxymethyl transferase (SHMT) reaction, though this does not explain the higher fraction of M+1 serine compared to its M+2.^37^ Further analysis is needed to understand the contributions of these metabolic pathways.

Elucidating the mechanisms underlying all of these metabolic changes is beyond the scope of the current work. We present the data here as a resource to offer a comprehensive overview of the metabolic insights in type 2 alveolar epithelial cells infected with different strains of Mtb. Our findings reveal how diverse Mtb strains predominantly adopt robust metabolic strategies with differential activities in the central carbon metabolism of host cells. However, further studies are needed to shed light on disease mechanisms regarding Mtb load and understand whether these changes stem from direct pathogen manipulation or host metabolic responses. This knowledge could guide deciphering the metabolic phenotypes using parallel ^13^C feeding (with positional labelled substrates), which will allow defining strategies to enhance immune responses and combat bacterial infections more effectively. Also, the current study elucidates the importance of deciphering the strain-specific host-pathogen metabolic interactions to strategies development of targeted therapies against drug-resistant Mtb infections.

### Limitations of the study

Our study has certain limitations. Firstly, these studies included A549 cells, and many such cell lines have undergone significant metabolic changes, potentially resembling those induced by bacterial pathogens, which could obscure pathogen-specific host responses. The primary human AEC2 cells are known to lose their phenotype and surfactant synthesis capacity during a standard culture *in vitro*.^38^ However, validation in primary alveolar epithelial cells will be useful. Secondly, we analyzed a relatively small number of clinical isolates (two drug-sensitive and two drug-resistant isolates). A larger number of clinical Mtb isolates with varying resistance profiles is needed to confirm these observations and demonstrate the universality of the results. Thirdly, further comprehensive and follow-up research is required for a more robust and convincing conclusion and to determine if the known effects are common to all drug-resistant Mtb isolates of different lineages.

## Acknowledgements

We acknowledge the CORE funding from the International Centre for Genetic Engineering and Biotechnology New Delhi to R.K.N., Fellowship support from the Department of Biotechnology, Government of India to S.A. and seed grant support from Indian Institute of Technology Mandi to SKM. Department of Biotechnology supported Tuberculosis Aerosol Challenge Facility (TACF) at ICGEB is kindly acknowledged. Sandeep Rai Kaushik, Sukanya Sahu and Shweta Chaudhary provided help during the experiments. We thank Ms Purnima Kumar for confocal microscopy image acquisition, Falak Pahwa for the PDIM experiment, Amit Kumar Mohapatra for his assistance with the antibiotic susceptibility assay of the clinical isolates, Haripriya for the heat-killed H37Rv electron microscopy experiment, and Manu Shree for discussions and feedback.

## Author contributions

S.A., S.K.M. and R.K.N. designed the experiments. R.K.N. supervised the study. S.A. performed the experiments and analyzed the data. S.A. and R.K.N. wrote the original draft. S.A., S.K.M., and R.K.N. reviewed and edited the manuscript. S.A., S.K.M., and R.K.N. contributed to data interpretation, and all authors approved the manuscript.

## Declaration of interests

The authors declare no competing interest.

## STAR★Methods

### Key resources table

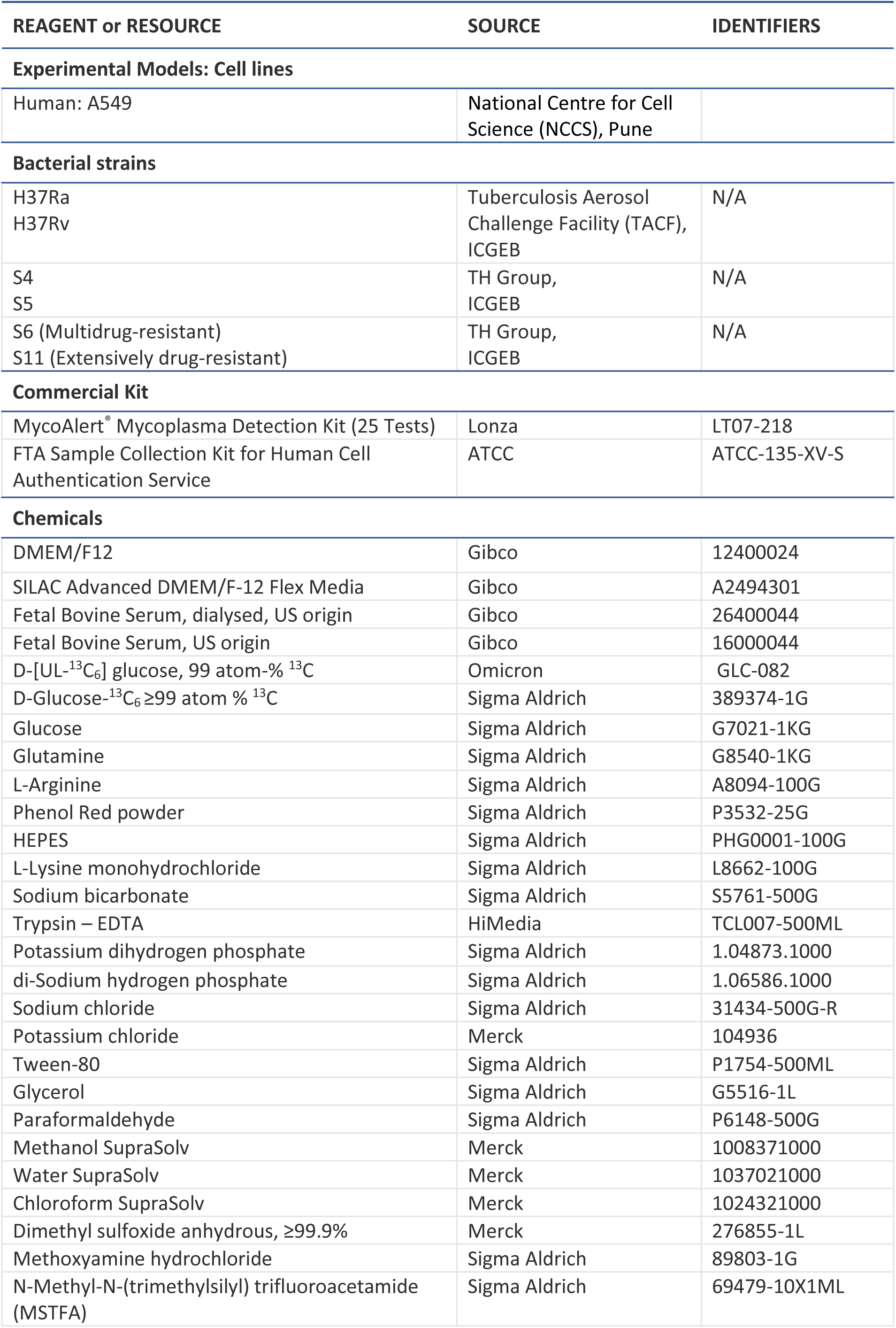
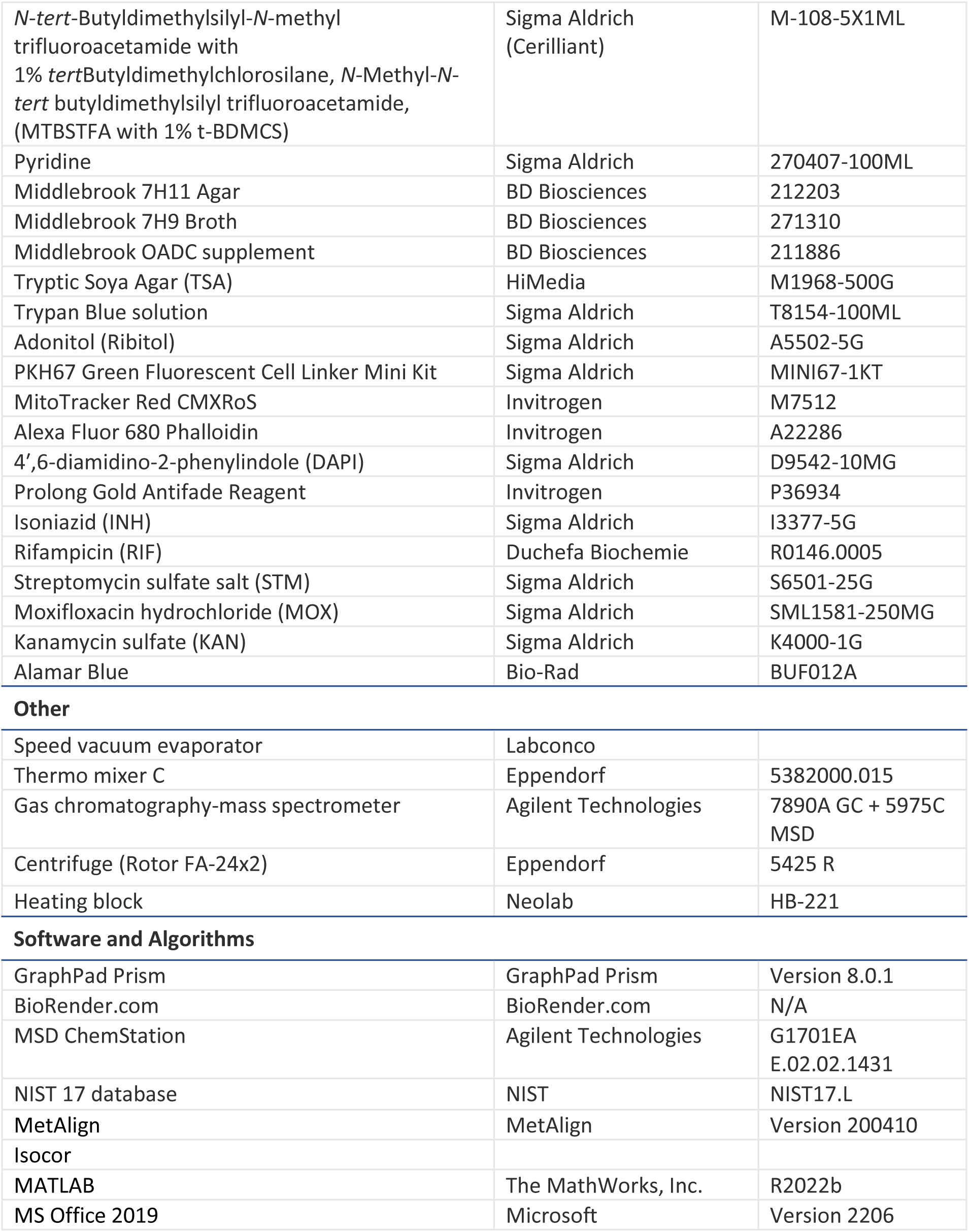

| REAGENT or RESOURCE | SOURCE | IDENTIFIERS |
| --- | --- | --- |
| **Experimental Models: Cell lines** |  |  |
| Human: A549 | National Centre for Cell Science (NCCS), Pune |  |
| **Bacterial strains** |  |  |
| H37Ra<br>H37Rv | Tuberculosis Aerosol Challenge Facility (TACF), ICGB | N/A |
| S4<br>S5 | TH Group, ICGB | N/A |
| S6 (Multidrug-resistant)<br>S11 (Extensively drug-resistant) | TH Group, ICGB | N/A |
| **Commercial Kit** |  |  |
| MycoAlert® Mycoplasma Detection Kit (25 Tests) | Lonza | LT07-218 |
| FTA Sample Collection Kit for Human Cell Authentication Service | ATCC | ATCC-135-XV-S |
| **Chemicals** |  |  |
| DMEM/F12 | Gibco | 12400024 |
| SILAC Advanced DMEM/F-12 Flex Media | Gibco | A2494301 |
| Fetal Bovine Serum, dialysed, US origin | Gibco | 26400044 |
| Fetal Bovine Serum, US origin | Gibco | 16000044 |
| D-[UL-<sup>13</sup>C<sub>6</sub>] glucose, 99 atom-% <sup>13</sup>C | Omicron | GLC-082 |
| D-Glucose-<sup>13</sup>C<sub>6</sub> ≥99 atom % <sup>13</sup>C | Sigma Aldrich | 389374-1G |
| Glucose | Sigma Aldrich | G7021-1KG |
| Glutamine | Sigma Aldrich | G8540-1KG |
| L-Arginine | Sigma Aldrich | A8094-100G |
| Phenol Red powder | Sigma Aldrich | P3532-25G |
| HEPES | Sigma Aldrich | PHG0001-100G |
| L-Lysine monohydrochloride | Sigma Aldrich | L8662-100G |
| Sodium bicarbonate | Sigma Aldrich | S5761-500G |
| Trypsin – EDTA | HiMedia | TCL007-500ML |
| Potassium dihydrogen phosphate | Sigma Aldrich | 1.04873.1000 |
| di-Sodium hydrogen phosphate | Sigma Aldrich | 1.06586.1000 |
| Sodium chloride | Sigma Aldrich | 31434-500G-R |
| Potassium chloride | Merck | 104936 |
| Tween-80 | Sigma Aldrich | P1754-500ML |
| Glycerol | Sigma Aldrich | G5516-1L |
| Paraformaldehyde | Sigma Aldrich | P6148-500G |
| Methanol SupraSolv | Merck | 1008371000 |
| Water SupraSolv | Merck | 1037021000 |
| Chloroform SupraSolv | Merck | 1024321000 |
| Dimethyl sulfoxide anhydrous, ≥99.9% | Merck | 276855-1L |
| Methoxyamine hydrochloride | Sigma Aldrich | 89803-1G |
| N-Methyl-N-(trimethylsilyl) trifluoroacetamide (MSTFA) | Sigma Aldrich | 69479-10X1ML |
| *N*-tert-Butyldimethylsilyl-*N*-methyl trifluoroacetamide with 1% *tert*Butyldimethylchlorosilane, *N*-Methyl-*N*-*tert* butyldimethylsilyl trifluoroacetamide, (MTBSTFA with 1% t-BDMCS) | Sigma Aldrich (Cerilliant) | M-108-5X1ML |
| Pyridine | Sigma Aldrich | 270407-100ML |
| Middlebrook 7H11 Agar | BD Biosciences | 212203 |
| Middlebrook 7H9 Broth | BD Biosciences | 271310 |
| Middlebrook OADC supplement | BD Biosciences | 211886 |
| Tryptic Soya Agar (TSA) | HiMedia | M1968-500G |
| Trypan Blue solution | Sigma Aldrich | T8154-100ML |
| Adonitol (Ribitol) | Sigma Aldrich | A5502-5G |
| PKH67 Green Fluorescent Cell Linker Mini Kit | Sigma Aldrich | MINI67-1KT |
| MitoTracker Red CMXRos | Invitrogen | M7512 |
| Alexa Fluor 680 Phalloidin | Invitrogen | A22286 |
| 4',6-diamidino-2-phenylindole (DAPI) | Sigma Aldrich | D9542-10MG |
| Prolong Gold Antifade Reagent | Invitrogen | P36934 |
| Isoniazid (INH) | Sigma Aldrich | I3377-5G |
| Rifampicin (RIF) | Duchefa Biochemie | R0146.0005 |
| Streptomycin sulfate salt (STM) | Sigma Aldrich | S6501-25G |
| Moxifloxacin hydrochloride (MOX) | Sigma Aldrich | SML1581-250MG |
| Kanamycin sulfate (KAN) | Sigma Aldrich | K4000-1G |
| Alamar Blue | Bio-Rad | BUF012A |
| Other |  |  |
| Speed vacuum evaporator | Labconco |  |
| Thermo mixer C | Eppendorf | 5382000.015 |
| Gas chromatography-mass spectrometer | Agilent Technologies | 7890A GC + 5975C MSD |
| Centrifuge (Rotor FA-24x2) | Eppendorf | 5425 R |
| Heating block | Neolab | HB-221 |
| Software and Algorithms |  |  |
| GraphPad Prism | GraphPad Prism | Version 8.0.1 |
| BioRender.com | BioRender.com | N/A |
| MSD ChemStation | Agilent Technologies | G1701EA<br>E.02.02.1431 |
| NIST 17 database | NIST | NIST17.L |
| MetAlign | MetAlign | Version 200410 |
| Isocor |  |  |
| MATLAB | The MathWorks, Inc. | R2022b |
| MS Office 2019 | Microsoft | Version 2206 |

### Experimental model and study participant details

#### Biosafety declaration

All the experimental procedures used in this study were approved by the Institutional Bio-Safety Committee (DBT Memorandum No. BT/BS/17/006/96-PID, dated 17/02/2016) and carried out at the Tuberculosis Aerosol Challenge Facility (TACF), a DBT-sponsored national facility of International Centre for Genetic Engineering and Biotechnology, New Delhi, following the Biosafety Label-III guidelines.

### Cell Culture

Type 2 alveolar epithelial adenocarcinoma cell line A549 was procured from the National Centre for Cell Science (NCCS), Pune (India) and authenticated by the American Type Culture Collection (ATCC). Before the experiments, the cells were confirmed to be mycoplasma-free using a MycoAlert detection kit (Lonza, LT07-218). A549 cells were maintained in DMEM/F12 (Gibco, 12400024) and 10% v/v heat-inactivated fetal bovine serum (US origin, Gibco, 16000044) supplemented with sodium bicarbonate (1.2 g/L) in a humidified incubator (Steri-Cycle CO_2_ incubator, Thermo Scientific) at 37 °C with 5% CO_2_. For ^13^C isotopic glucose labelling experiments (post-infection), cells were maintained in SILAC Advanced DMEM/F-12 Flex Media (Gibco, A2494301) and 5% v/v heat-inactivated dialysed fetal bovine serum (US origin, Gibco, 26400044) supplemented with glucose (17.50 mM, U-^13^C_6_/U-^12^C_6_), L-Glutamine (2.50 mM), L-Arginine (0.69 mM), L-Lysine monohydrochloride (0.49 mM), HEPES (15.01 mM) and Phenol Red (0.02 mM) in a humidified 37 °C incubator with 5% CO_2_. The medium was replaced every two to three days with fresh DMEM-F12. Cells were passaged when 75-90% confluent. For subculture, the culture medium was removed from the flasks, and the cell layer was washed twice with a warm serum-free DMEM/F12 medium. Trypsin EDTA was added, and the cells were placed back into the incubator. After 5 min of incubation, the culture flasks were gently tapped to dislodge cells from the surface. A complete DMEM/F12 medium (DMEM/F12 supplemented with FBS) at a volume ratio of 1:2 (trypsin: medium) was added to the flask, and the cell suspension was transferred to a conical centrifuge tube. The cells were centrifuged, resuspended in a single-cell suspension in fresh medium, and seeded in tissue culture flasks at a ratio of 1:3. Cells between passages 5-8 were used for all experiments. Cell morphology was monitored in all cultures using a light microscope (Nikon).

### Bacteria culture

All laboratory Mtb strains (H37Rv and H37Ra) and clinical isolates (Drug-sensitive isolates S4 and S5, Drug-resistant isolates S6 and S11) were grown in Middlebrook 7H9 broth supplemented with glycerol (0.2% v/v), Tween-80 (0.05% v/v) and OADC (Oleic acid, Bovine Albumin, Dextrose, and Catalase; 10% v/v). Mtb cultures (10 mL) were incubated in 50 mL conical tubes at 37 °C under constant rotation at 180 rpm. Growth was monitored by measuring the optical density (OD) at 600 nm. Cultures were harvested in the mid-logarithmic phase (OD_600nm_ = 0.6), stocks prepared in 50% glycerol in 1 mL aliquots and stored at −80 °C. Before the experiment, aliquots were thawed and expanded in Middlebrook 7H9 broth. Aliquots of bacterial cultures were plated on tryptic soy agar (TSA) to check for contamination. All Mtb strains were tested for PDIM (Phthiocerol dimycocerosates) positivity status using thin-layer chromatography (TLC) of lipid extracts from cultures before performing the infection experiments.

### PDIM positivity test

Mtb cultures were grown to the mid-logarithmic phase (O.D._600nm_ = 0.6). The cultures were pelleted, and a mixture of Phenol: Chloroform: Water (10:10:3, 10 mL) was added to each sample and incubated overnight (∼16 h) in a shaker incubator at 37 °C and centrifuged at 3,500 *g* for 10 min at room temperature. The supernatant was filtered using a 0.2 µm nylon membrane syringe filter. The filtered supernatant was vacuum-dried in 2 mL microcentrifuge tubes. Lipid samples were resuspended in methanol (100 µL), and aliquots were spotted onto thin layer chromatography (TLC) plates (250 µm silica gel 60 F_254_ plates). Preparative TLCs were resolved using petroleum ether/ethyl acetate (98:2, v/v) as the solvent. TLC was developed in an iodine chamber to detect the PDIM bands (Figure S1).

### Heat-killed Mtb preparation

A single-cell suspension of the H37Rv strain was heated at 95 °C in a water bath for 15 minutes, cooled to room temperature, and resuspended in DMEM/F-12 Flex medium for infection experiments.^39^ Heat-killed H37Rv Mtb strain was characterised using electron microscopy (Figure S3).

### Antibiotic susceptibility test of clinical Mycobacterial isolates

Analysis and antibiotic sensitivity characterisation of Mtb culture, isolated from sputum samples of tuberculosis patients, was performed by a NABL (National Accreditation Board for Testing and Calibration Laboratories) accredited laboratory. Antimicrobial susceptibility testing (AST) of selected clinical Mtb isolates was reperformed in the laboratory using the Alamar Blue assay. Harvested bacterial cells in the log growth phase and cell counts were determined by measuring optical density (OD) at 600 nm. An aliquot of the Mtb culture (∼6×10^6^ bacteria) was transferred to a microcentrifuge tube and centrifuged at 3,000 *g* for 5 min. The supernatant was discarded, and the pellet was resuspended in fresh 7H9 medium supplemented with OADC (10%, 500 µL). A single-cell suspension of the Mtb culture was prepared, and the cell count was adjusted to 1×10^5^ bacteria per well. Bacterial cells were treated with various concentrations of anti-tuberculosis drugs (Isoniazid, Rifampicin, Streptomycin, Kanamycin and Moxifloxacin) in 96-well plates. All outer perimeter wells were filled with sterile water to prevent dehydration of the medium in the experimental wells. In the control wells, only bacteria or media were added. The bacterial cells were incubated with the drugs for 5 days. AlamarBlue (20 µL; 10% of the total volume of 200 µL per well) was added aseptically to the wells. The plates were incubated with alamarBlue dye for 24 h and the colour change from blue to pink was monitored at 12 and 24 h.

### Cell Infection

Mtb cultures in the mid-logarithmic phase (O.D._600 nm_ = 0.6) were centrifuged at 4,000 *g* for 5 minutes at RT, and the pellets were resuspended in serum-free DMEM/F12 culture media. Mycobacteria were passed through a 23-gauge needle, followed by a 26-gauge needle 10 times each, to prepare a single-cell suspension. The bacterial culture at 0.6 O.D., corresponds to 100 million bacteria per mL at a wavelength of 600 nm.^40^ The required number of bacteria was added to the seeded cells in a complete medium at a multiplicity of infection (MOI) of 1:10 (cell: bacteria). At 8 h post-infection, the media was removed, the cells were washed three times with sterile phosphate-buffered saline (PBS) to remove extracellular bacilli, and fresh complete media was added. Cells were placed in a humidified incubator at 37 °C and 5% CO_2_ for the specified times. This infection protocol was followed for all subsequent experiments.

### Cell proliferation and viability assay

A549 cells were cultured in 6-well plates at 0.6 million per mL density, allowed to adhere for 24-36 h, infected with Mtb at MOI 1:10, and harvested at designated time points for manual counting using the trypan blue reagent. The cells were diluted at 1:10 or 1:20 with trypan blue, and 10 μL cell suspension was added to a Neubauer haemocytometer and counted under a light microscope (Nikon). The cell count/mL was calculated as follows:

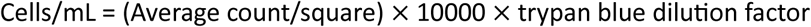

### Intracellular bacteria viability assay (Colony forming unit or CFU)

A colony-forming unit (CFU) assay using the drop-plate method was performed to determine intracellular bacterial counts. The extracellular medium was removed, and the infected cells were washed three times with sterile PBS (1×) and lysed with SDS (0.06%, prepared in PBS). Diluted cell lysates (1,000 times dilution in 7H9 media) were plated onto 7H11 plates in duplicate and incubated at 37 °C. CFU counts were enumerated after 21 and 42 days of incubation for Mtb laboratory strains and clinical isolates, respectively, and were expressed as CFU/number of viable cells. The remaining cells were incubated for further time points before lysis, dilution, and plating.

### Confocal microscopy

Mtb H37Rv bacteria were stained with PKH67 (Sigma Aldrich, MINI67-1KT), a green fluorescent lipophilic dye, following the manufacturer’s protocol. Briefly, Mtb cultures (O.D._600 nm_ = 0.6) were centrifuged at 4,000 rpm for 5 minutes, and the bacterial pellet was washed with PBS. Each pellet was resuspended in diluent (250 µL). Dye was added to stain the bacteria corresponding to 2 µL dye per 1,000 million bacteria and incubated for 7 min at room temperature (RT) in the dark. FBS (500 µL, twice the volume of the diluent) was added, and the mixture was incubated for 2 min at RT in the dark. The bacteria were washed twice with PBS and resuspended in serum-free DMEM/F12 to prepare single-cell suspensions.

A549 cells were seeded at 0.06 million/mL density onto 18 mm coverslips in 12-well plates and allowed to adhere for 24-36 h at 37 °C in a 5% CO_2_ humidified incubator. The cells were infected with fluorescent dye labelled Mtb. Cells were washed with PBS, and mitochondria were stained with MitoTracker® red (200 nM) for 30 min in a humidified 5% CO_2_ incubator at 37 °C. After washing three times with PBS, the cells were fixed with paraformaldehyde (PFA, 2%) for 30 min at room temperature in the dark. The fixed cells were washed thrice with PBS and permeabilised with 0.1% Triton-X-100 for 5 min. It was followed by washing the cells thrice with PBS and blocking them with 1% BSA for 20 min. For actin staining, Alexa Fluor^TM^ 680 phalloidin (∼1 unit/sample) was added to the cells and incubated for 20 min at RT. The cells were washed thrice with PBS, and the cell nuclei were stained with 4′,6-diamidino-2-phenylindole (DAPI, 1 µg/ml) for 15 min at RT in the dark. The cells were given a final wash three times with PBS and mounted onto glass slides using the ProLong Gold antifade reagent. All solutions were prepared in PBS. Images were acquired using NIS-Elements AR analysis software (version 4.0, Life Technologies, USA) using a Nikon A1R Laser Scanning Confocal Microscope equipped with a Nikon Plan Apo VC ×60, NA 0.75, and Plan Apo VC ×100 oil with 1.40 numerical aperture (NA), an oil immersion objective lens. They were visualised using DAPI (Ex/Em-358/461 nm), PE/AlexaFluor680 (Ex/Em-679/702 nm), TRITC (Ex/Em-579/599 nm), FITC (Ex/Em-490/502 nm) and TD (bright field) filters and NIS-Elements AR analysis software (version 4.0, Life Technologies, USA).

### Metabolite extraction

A549 cells were seeded at the required density in 6-well plates, infected with Mtb, and incubated at 37 °C in a humidified incubator with 5% CO_2_. At 8 h post-infection, the media was removed, the cells were washed three times with sterile PBS, and fresh complete media (supplemented with U-^12^C_6_ /U-^13^C_6_ glucose, 17.50 mM) was added. This step was considered to be time-point 0 (time zero). The cells were then placed in a humidified incubator at 37 °C and 5% CO_2_. Cell culture media (ccm) were collected at designated time points, filtered using syringe filters (0.22 µm, PES membrane), and stored at −80 °C until further analysis. The cells were washed with ice-cold saline (0.9% NaCl) and quenched with pre-chilled methanol (800 μL), followed by the addition of ice-cold Milli-Q water (800 μL). The cells were scraped and collected in cold 15 mL conical centrifuge tubes. A pre-chilled chloroform (1.6 ml) was added to the tubes and vortexed at 4 °C for 30 min. The samples were centrifuged at 10,000 *g* for 10 min at 4 °C. The upper aqueous phase (methanolic) and lower organic phase (chloroform) were separated and stored at −80 °C until further analysis. The junction layers of the aqueous and organic phases comprising proteins were collected in screw-capped tubes and selected for amino acid analysis.

### Protein hydrolysis

The collected protein samples (n=450) were acid-hydrolysed by resuspending in 100 μL of 6M HCl. The tubes were partially sealed and placed inside a fume hood cabinet in a heating block for 12-14 h at 100 °C.^41^ The hydrolysates were centrifuged at 13,000 *g* for 10 min to separate water-soluble amino acids from the insoluble pellets. The supernatant from each sample was collected in a fresh tube and vacuum-dried at 22 °C for approximately 6-8 h in a speed vacuum evaporator. The dried hydrolysates were subjected to derivatisation for Gas chromatography-mass spectrometry (GC-MS).

### Chemical derivatisation of proteinogenic amino acids

Equal volumes of commercially available amino acids (n=22) were pooled to prepare quality control (QC) samples. Dried protein hydrolysates were subjected to *tert*-butyldimethylsilyl (TBDM) derivatisation. Pyridine (25 μL) was added to each dried sample, and the tubes were incubated at 37 °C for 30 min on a thermomixer set at 900 rpm. Then, MTBSTFA+1% t-BDMCS (35 μL, *N*-*tert*-Butyldimethylsilyl-*N*-methyltrifluoroacetamide + % *tert*-Butyldimethylchlorosilane, *N*-Methyl-*N*-*tert*-butyldimethylsilyltrifluoroacetamide) was added followed by incubation at 60 °C for 30 minutes in thermomixer at 900 rpm. All processed samples were centrifuged (10 min; 16,000 *g* at RT). The supernatant was transferred to 200 µL glass vial inserts in a GC vial (2 mL capacity) and subjected to GC-MS data acquisition.

### GC-MS data acquisition

GC-MS data acquisition of all samples was performed using a 7890 Gas Chromatograph and a 5975C quadrupole mass spectrometer (Agilent Technologies, Santa Clara, CA, USA). All samples (n=450) with separate sets of QCs were run in a single batch. Samples were randomized using an online tool (www.randomizer.org) and processed in a batch of 20 samples per day with 2 QC samples to run at the beginning and end of the sequence. Derivatised samples (1 μL) were injected into the GC column (Restek, RTX-5, 30 m × 250 µm × 0.25 µm) in a purged splitless mode.

For the TBDMS-derivatised samples, helium gas was used as the carrier gas at a constant flow rate of 1.3 mL/min with a solvent delay of 6 min. The oven program was set at 120 °C for 5 min, ramped to 270 °C at 4 °C/min, held for 3 min, ramped to 320 °C at 20 °C/min, and held for 1 min. The EI ion source temperature was set at 230 °C, the quadrupole temperature was set at 150 °C, and data were acquired in the full scan mode from 50 to 600 m/z for a total run time of 49 min. Ions were generated using a 70 eV electron beam in electron ionisation mode. An Agilent ChemStation software (version G1701EA E.02.02.1431) was used to control the data acquisition parameters during GC separation and mass spectra acquisition during all the sample runs. Data acquisition of samples was completed within 24-48 hours of derivatisation.

### GC-MS data analysis

The GC-MS spectra were baseline-corrected using MetAlign software^42^ to estimate the accurate mass isotopomer distributions (MIDs) for all amino acids. The peaks of the baselined spectra were identified using ChemStation software (version G1701EA E.02.02.1431) based on the m/z of different fragments, their retention times and hits against the NIST17 (National Institute of Standards and Technology, Maryland) Library and confirmed their identity by running commercial standards (Figure S2). The incorporated natural abundance of the ^13^C isotopes contributes to the measured MIDs. For accurate estimation of MIDs were corrected for natural isotope correction using Isocor software.^43^ The derivatisation and metabolite formula required for Isocor was calculated manually for all the metabolite fragments. The final corrected MIDs were used to calculate the average ^13^C abundance for each fragment. The amino acid fragments were validated by comparing the corrected average ^13^C abundance in labelled samples with fragments derived from unlabelled samples (in which the percentage of ^13^C is assumed to be less than the natural abundance, i.e., ≤1.13%). Thus, amino acid fragments derived from unlabelled samples with an average ^13^C ≤2% were assigned as valid.

Before statistical analysis, manual data curation was carried out to remove outliers. MetaboAnalyst 5.0 online tool was used for principal component analysis (PCA) and partial least square-discriminate analysis (PLS-DA). Missing values were replaced by 1/5 of the minimum positive values of their corresponding variables. Log transformation of data with auto-scaling was selected to obtain a near-normal distribution. Analytes with a Variable Importance projection (VIP) score >1.0 from the PLS-DA model were selected as important features to classify the groups.

### Statistical Analysis

Data compilation was performed using Microsoft Excel (2019). Plots were prepared, and statistical significance was measured using GraphPad Prism (version 8.0.1) with the following tests: Multiple t-tests, one-way ANOVA, and two-way ANOVA with multiple comparison analyses, as indicated in the respective figure legend. Significant comparisons were indicated as follows: p-values: * ≤ 0.05, ** <0.005, *** <0.0005, **** <0.0001 at 95% confidence interval. Error bars represent the standard deviation between replicates. The illustrations were created using BioRender (biorender.com).

### Data Availability

The metabolite data presented in this study are available at: https://data.mendeley.com/datasets/y6vxg79h5r/1

## Notes

https://data.mendeley.com/datasets/y6vxg79h5r/1

### Competing Interest Statement

The authors have declared no competing interest.

