## Supplementary data for "Isotopic tracing in *Mycobacterium tuberculosis*-infected alveolar epithelial cells indicates metabolic specificity to their virulence and drug-resistance status"

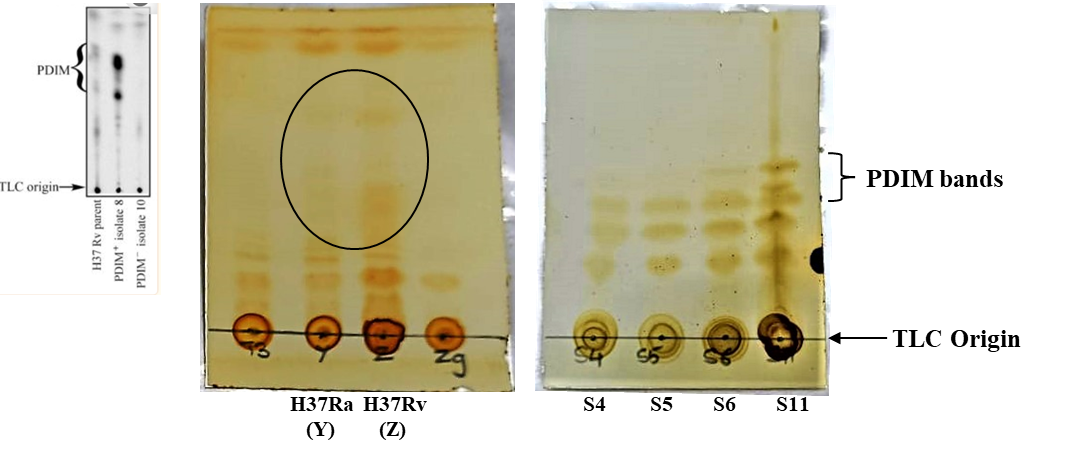

**Supplementary Figure 1:** Thin layer chromatography (TLC) plates developed in an iodine chamber showing the Phthiocerol dimycocerosates (PDIM) bands harvested from different *Mycobacterium tuberculosis* laboratory strains (H37Ra, H37Rv) and drug-sensitive and resistant clinical isolates.

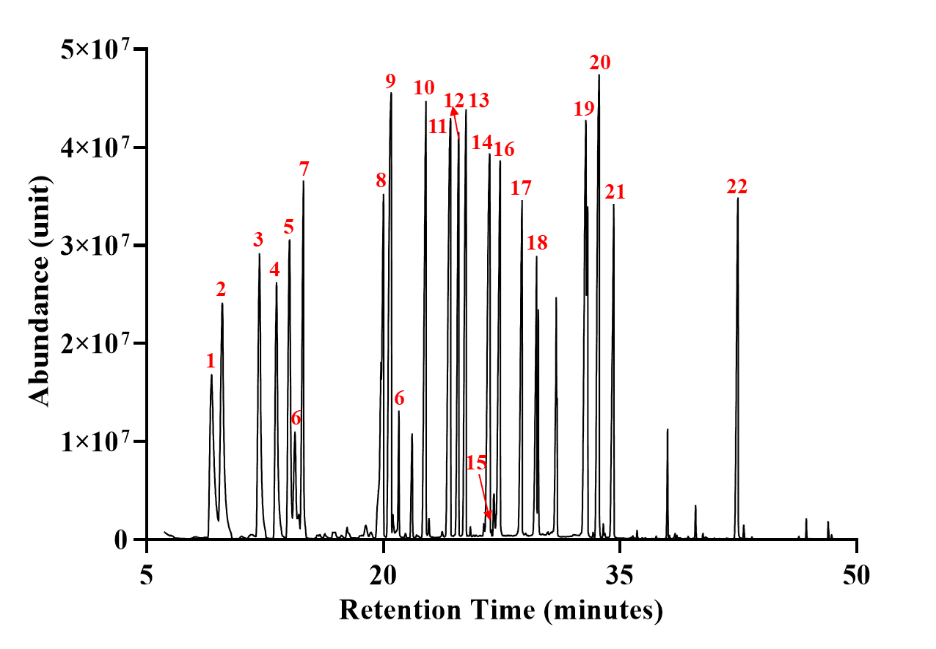

**Supplementary Figure 2:** **Representative total ion chromatogram (TIC) of GC-MS showing the distribution of TBDMS derivatized amino acids commercial standards (n=22)**. The amino acids are distributed based on their retention time - **1)** L-Alanine; **2)** L-Glycine; **3)** L-Valine; **4)** L-Leucine; **5)** L-Isoleucine; **6)** L-Threonine; **7)** L-Proline; **8)** L-Methionine; **9)** L-Serine; **10)** L-Phenylalanine; **11)** Aspartic acid; **12)** L-Hydroxyproline; **13)** L-Cysteine; **14)** L-Glutamic acid; **15)** L-Arginine; **16)** L-Asparagine; **17)** L-Lysine; **18)** L-Glutamine; **19)** L-Histidine; **20)** L-Tyrosine; **21)** L-Tryptophan; and **22)** L-Cystine.

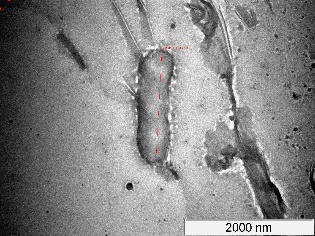

**Supplementary Figure 3:** Electron micrograph showing the characteristics of heat-killed *Mycobacterium tuberculosis* H37Rv strain.

**Supplementary Table 1:** Clinical *Mycobacterium tuberculosis* isolates used in this study.

| **Clinical Mtb**  **isolates** | **Classification** | **Drugs** | | | | |
| --- | --- | --- | --- | --- | --- | --- |
|  |  | **STM** | **INH** | **RIF** | **MOX** | **KAN** |
| S4 | Drug-sensitive | S | S | S | S | S |
| S5 |  | S | S | S | S | S |
| S6 | Drug-resistant (MDR) | R | R | R | S | S |
| S11 | Drug-resistant (XDR) | R | R | S | R | S |

**S - sensitive; R - resistant**

**Supplementary Table 2**: Average ^13^C abundance (in %) in the proteinogenic amino acid fragments of A549 cells cultured in DMEM/F12 medium containing either [U-^13^C_6_] or [U-^12^C_6_] glucose and infected with H37Rv or H37Ra or heat-killed H37Rv with uninfected cells as control (replicates=5). Std. dev. - Standard deviation. Related to Figure 3.

**a. Uninfected A549 cells**

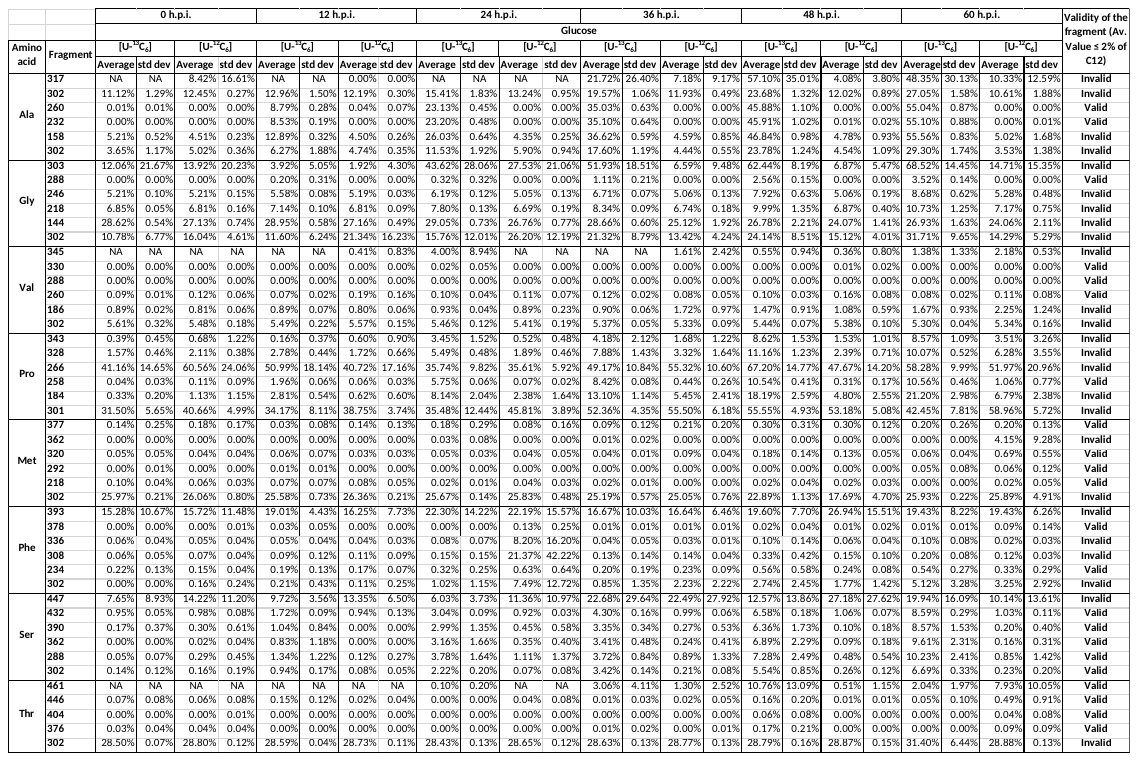

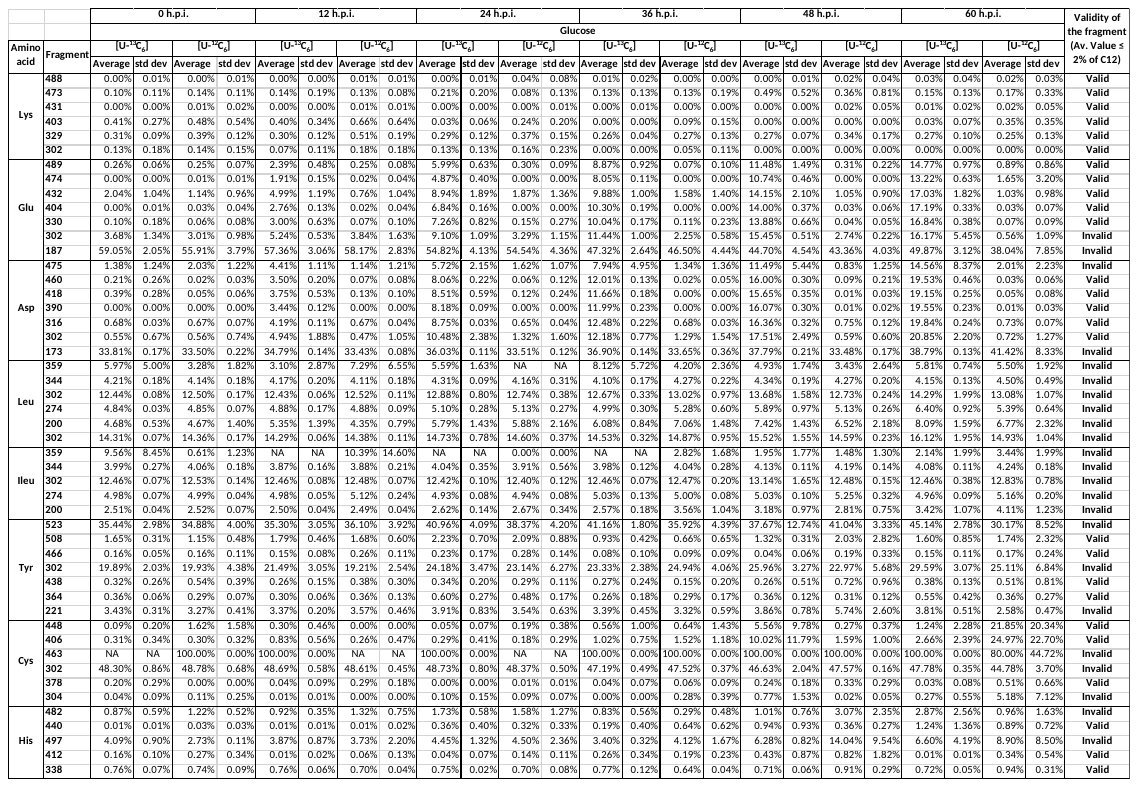

**b.** **Heat-killed Mtb H37Rv-infected A549 cells**

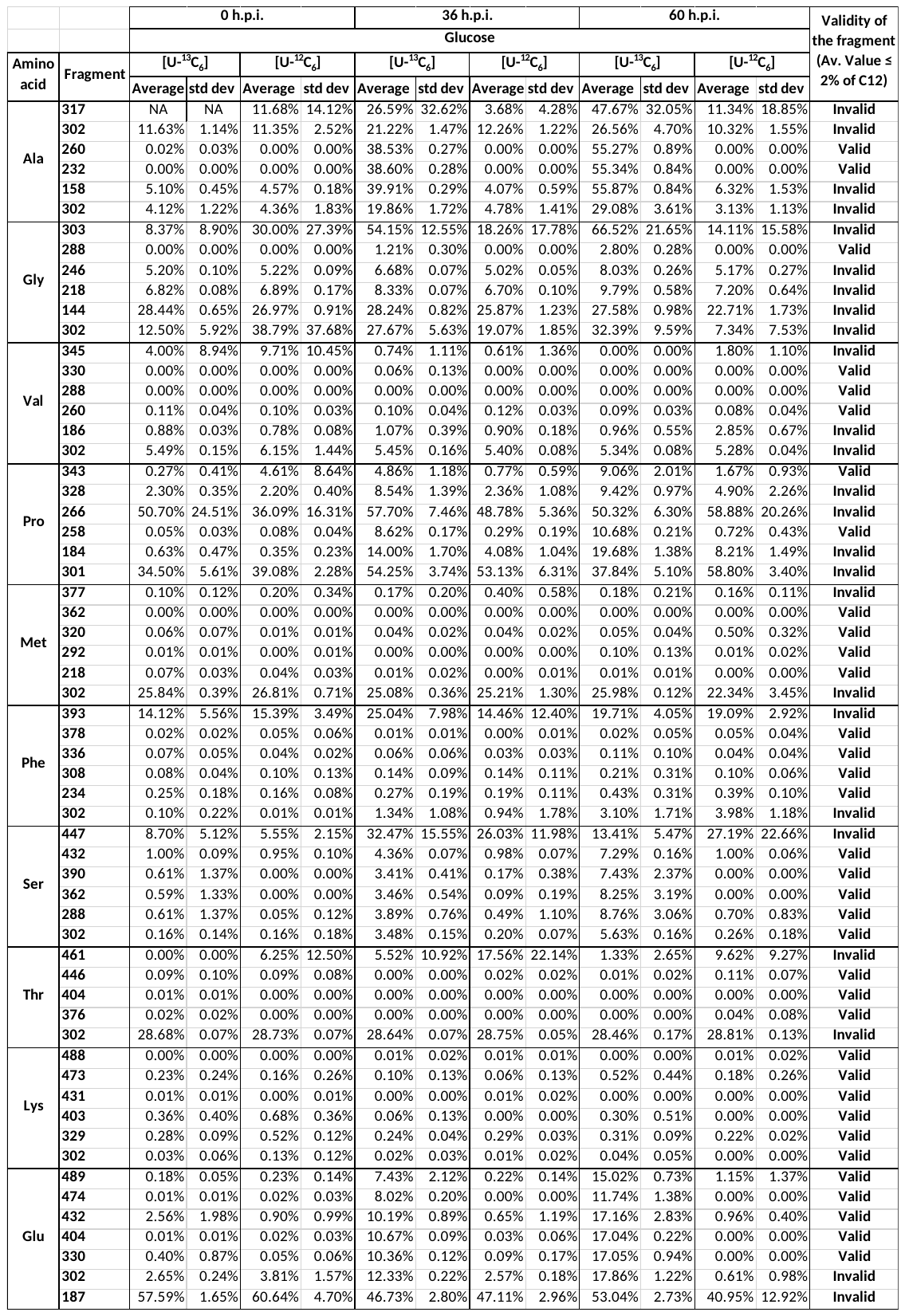

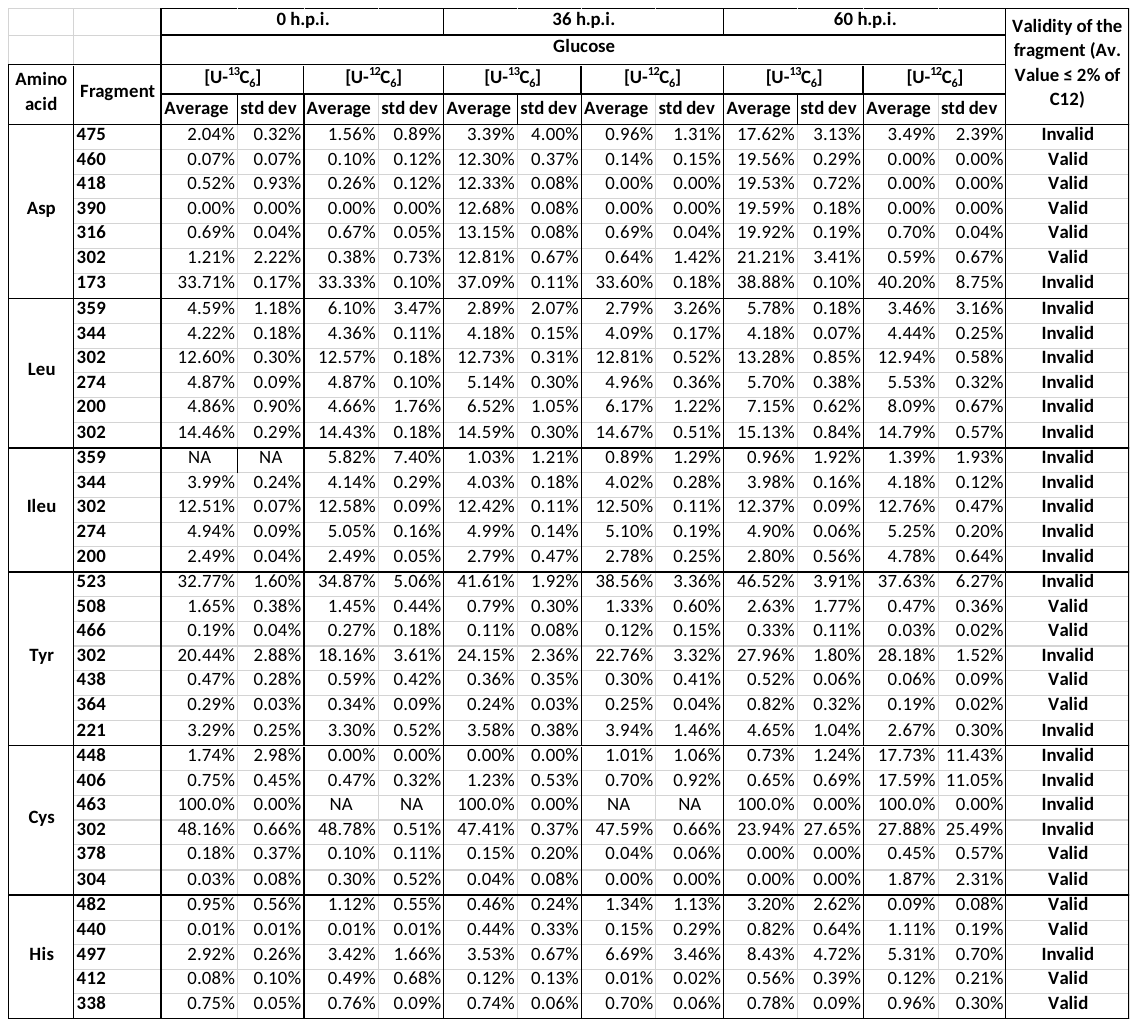

**c.** **H37Rv Mtb-infected A549 cells**

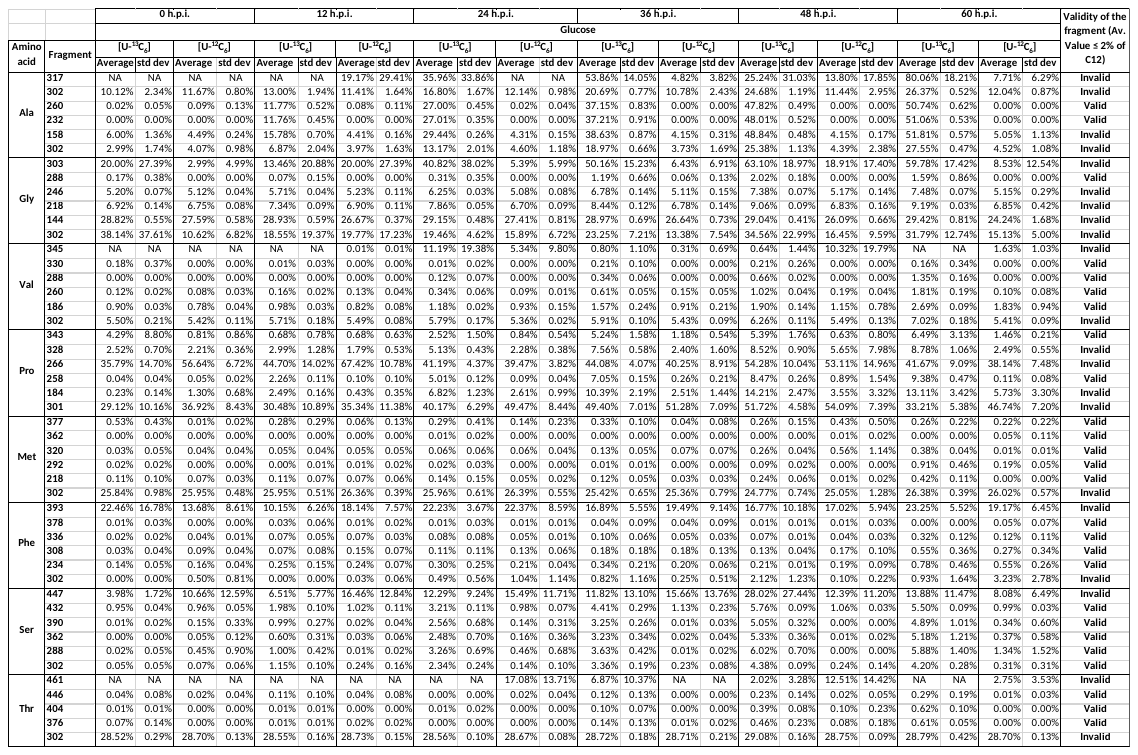

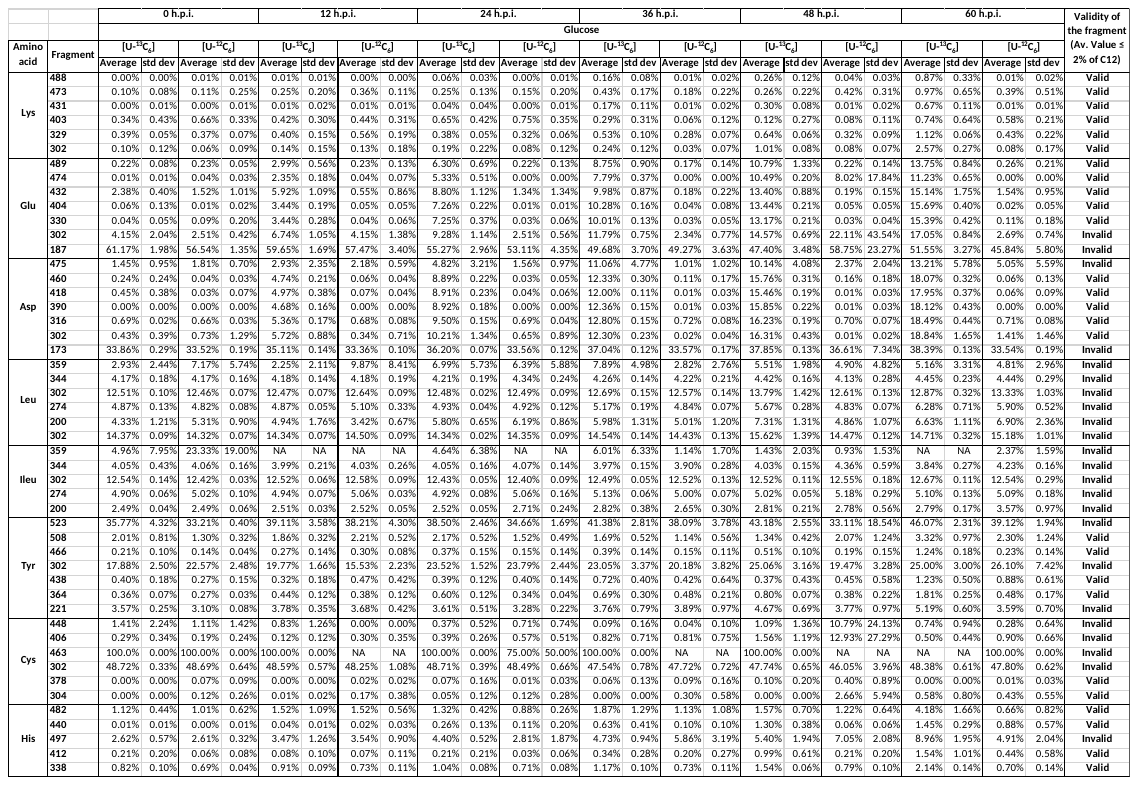
**d. H37Ra Mtb-infected A549 cells**

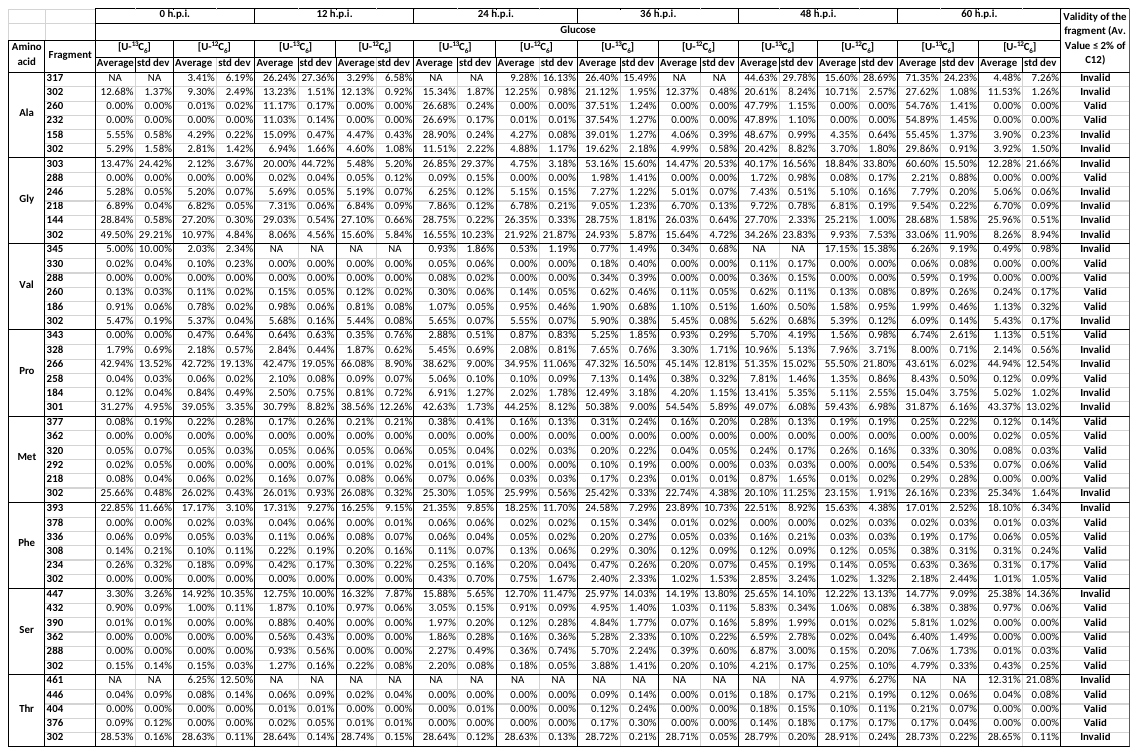

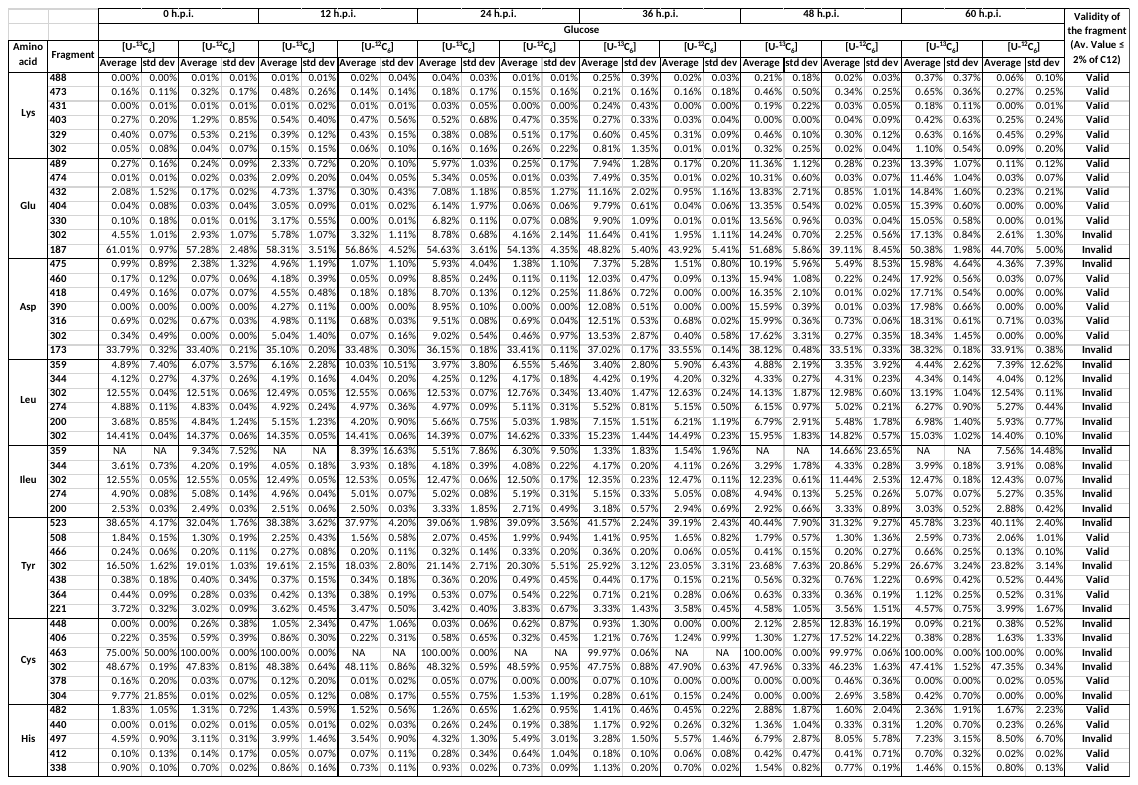
**Supplementary Table 3:** Mass isotopomer distribution (MID) of the proteinogenic acid fragments of A549 cells at 60 h.p.i. cultured in DMEM/F12 medium containing either [U-^13^C_6_] or [U-^12^C_6_] glucose and infected with H37Rv or H37Ra or heat-killed H37Rv with uninfected cells as control (replicates=5). Std. dev. - Standard deviation. Related to Figure 4.

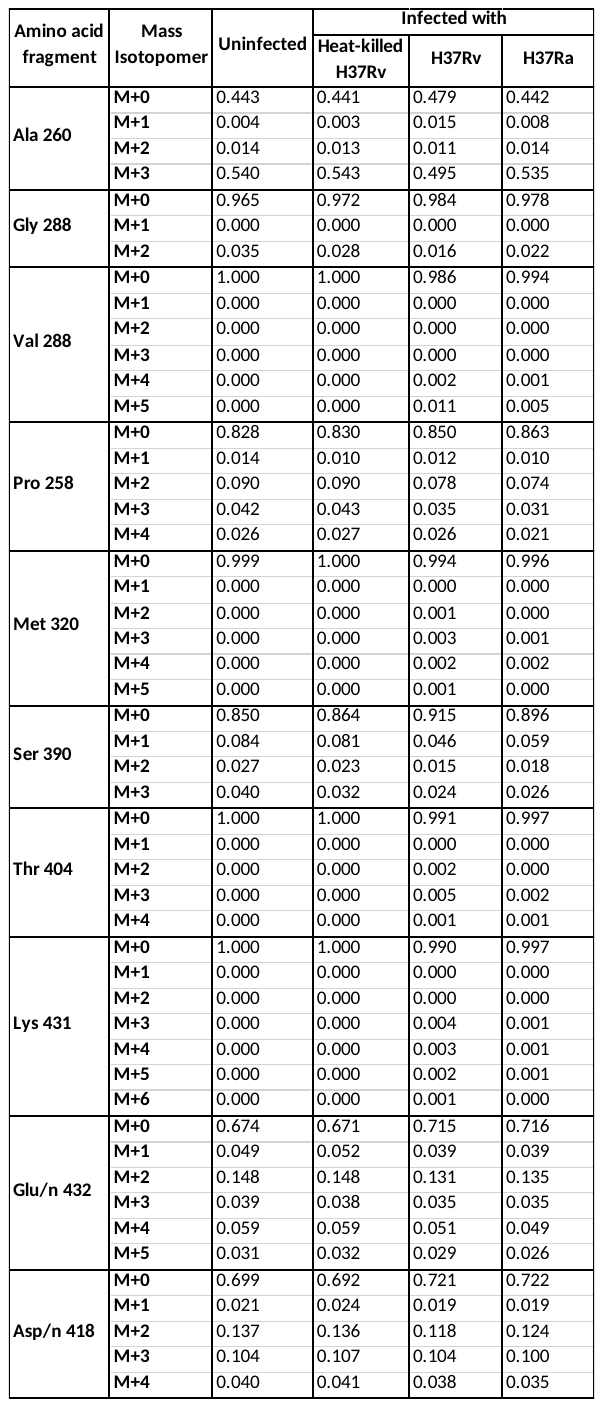

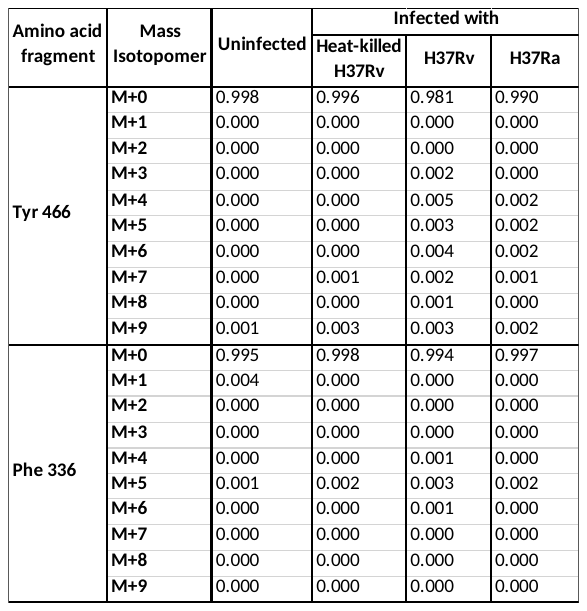

**Supplementary Table 4:** Average ^13^C abundance (in %) in the proteinogenic amino acid fragments of A549 cells cultured in DMEM/F12 medium containing either [U-^13^C_6_] or [U-^12^C_6_] glucose and infected with clinical *Mycobacterium tuberculosis* (Mtb) isolates (S4, S5, S6 and S11) and reference laboratory strain H37Rv with uninfected cells as control (replicates=5). Std dev: Standard deviation. Related to Figure 7.

**a. Uninfected A549 cells**

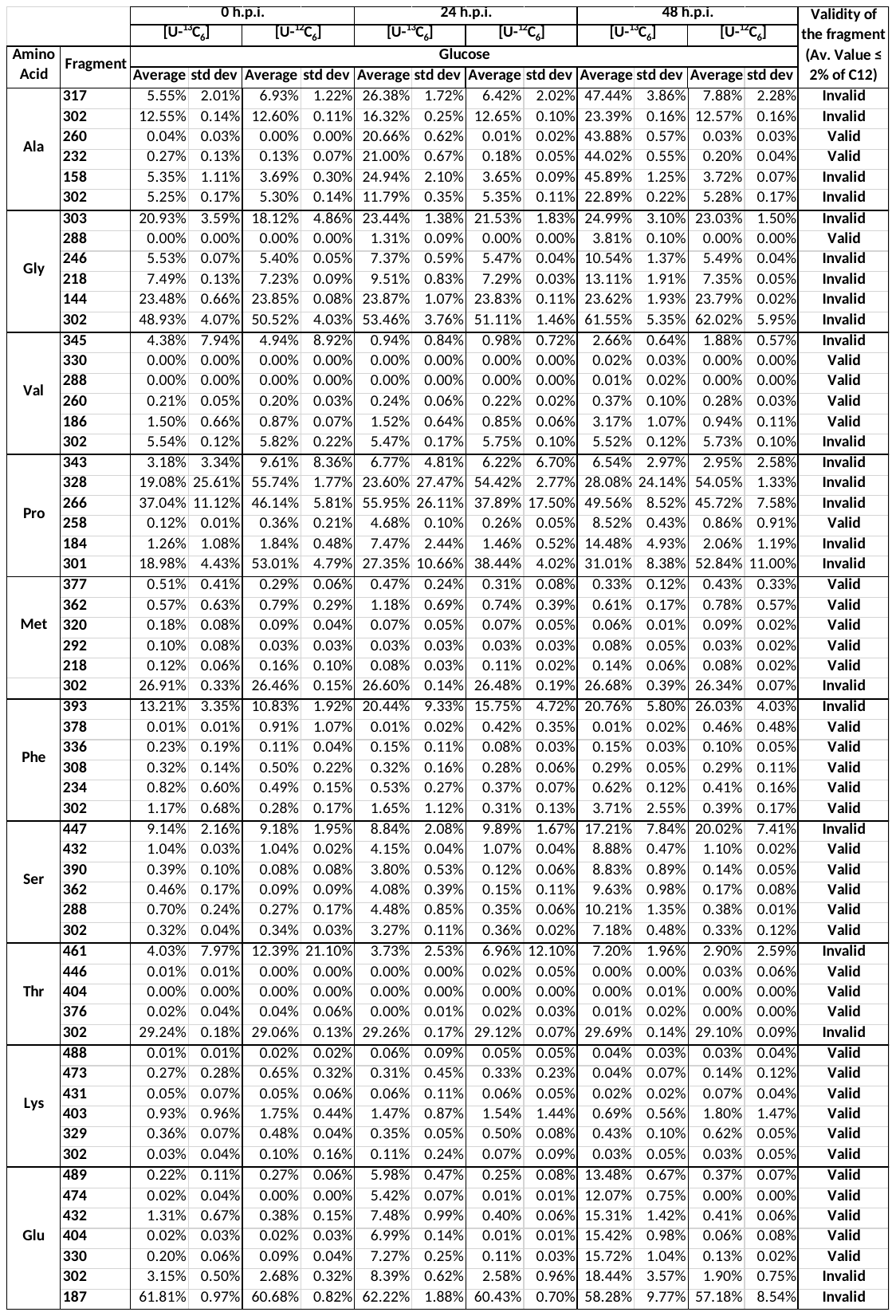

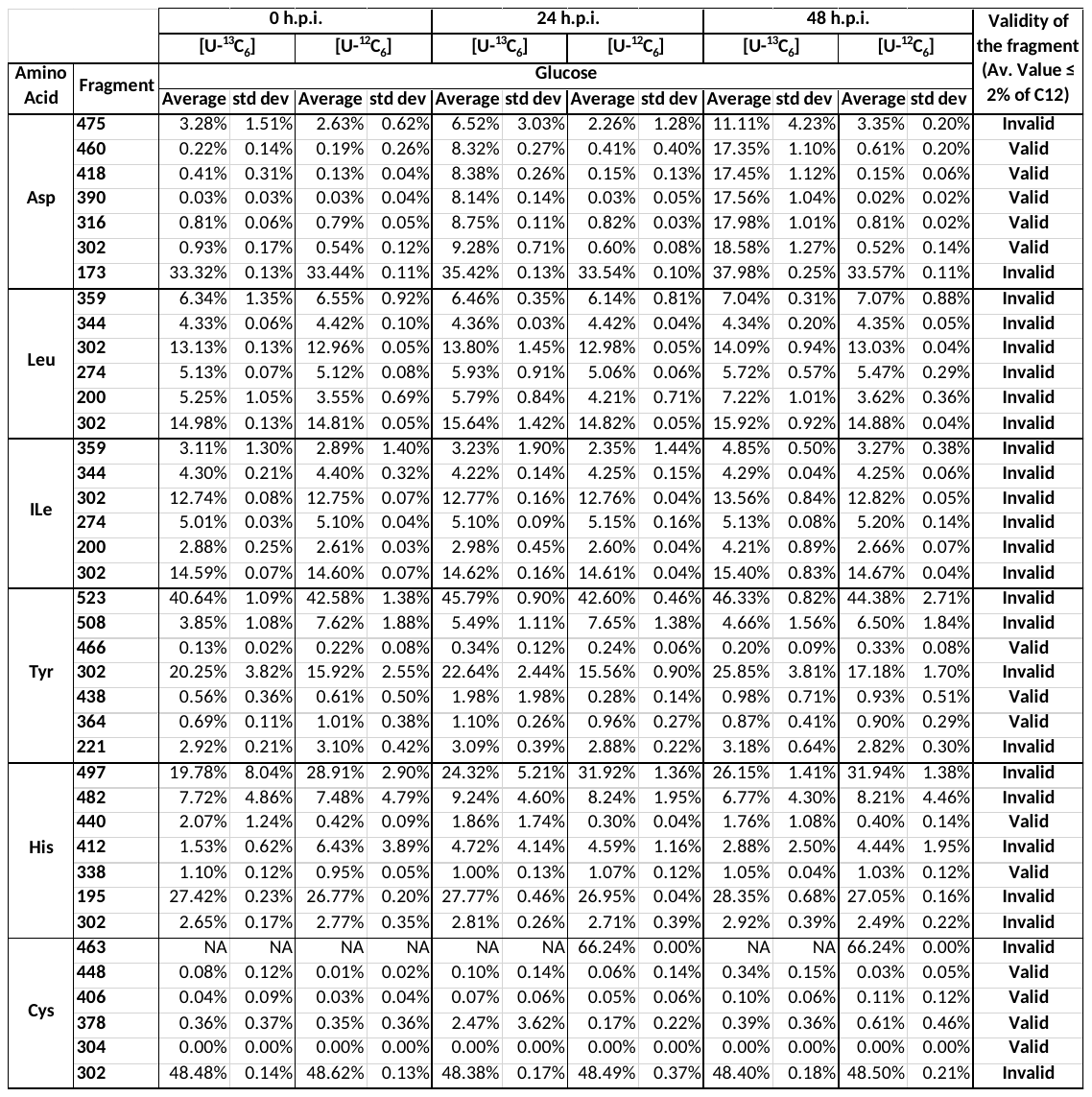

**b. Mtb H37Rv-infected A549 cells**

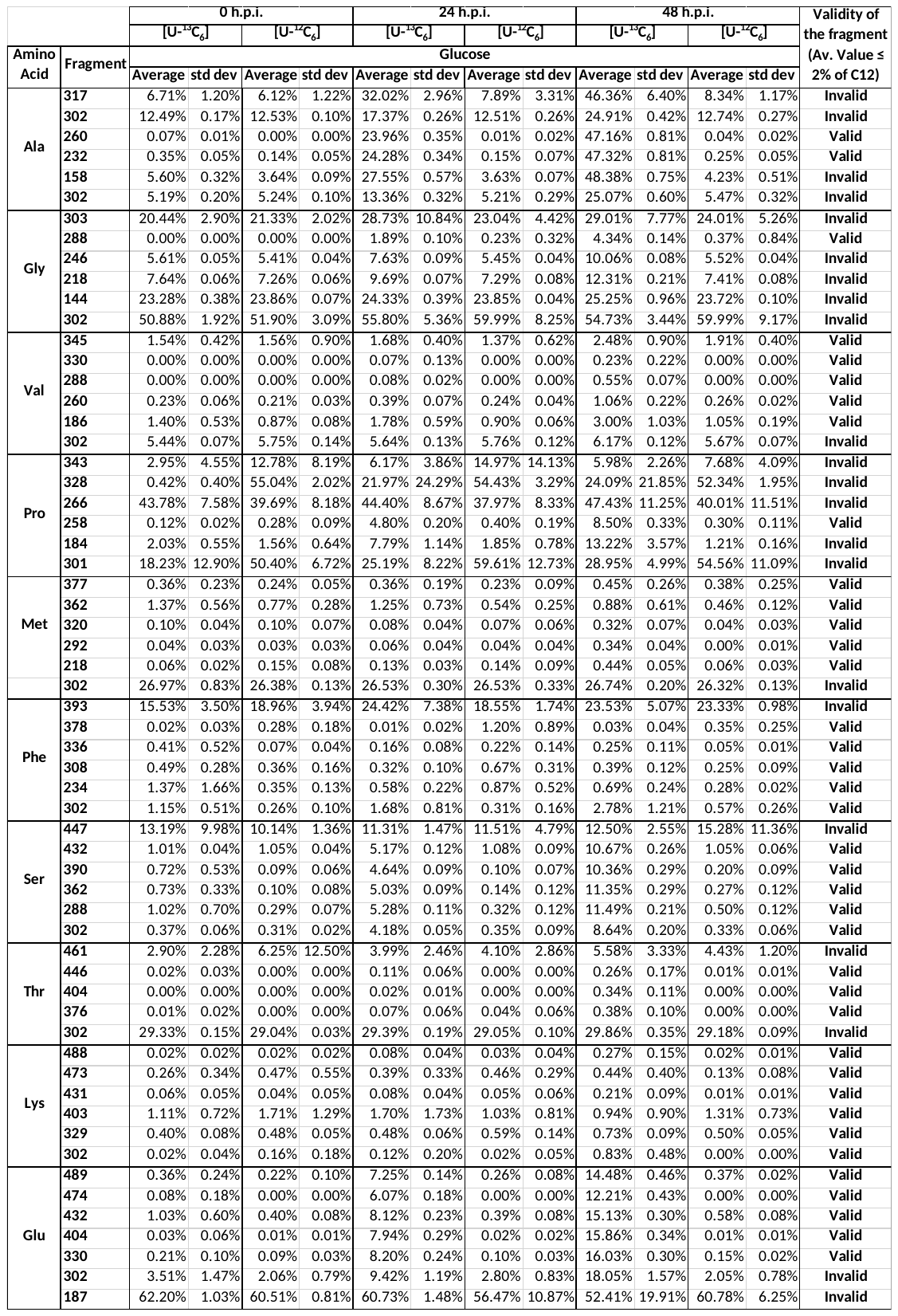

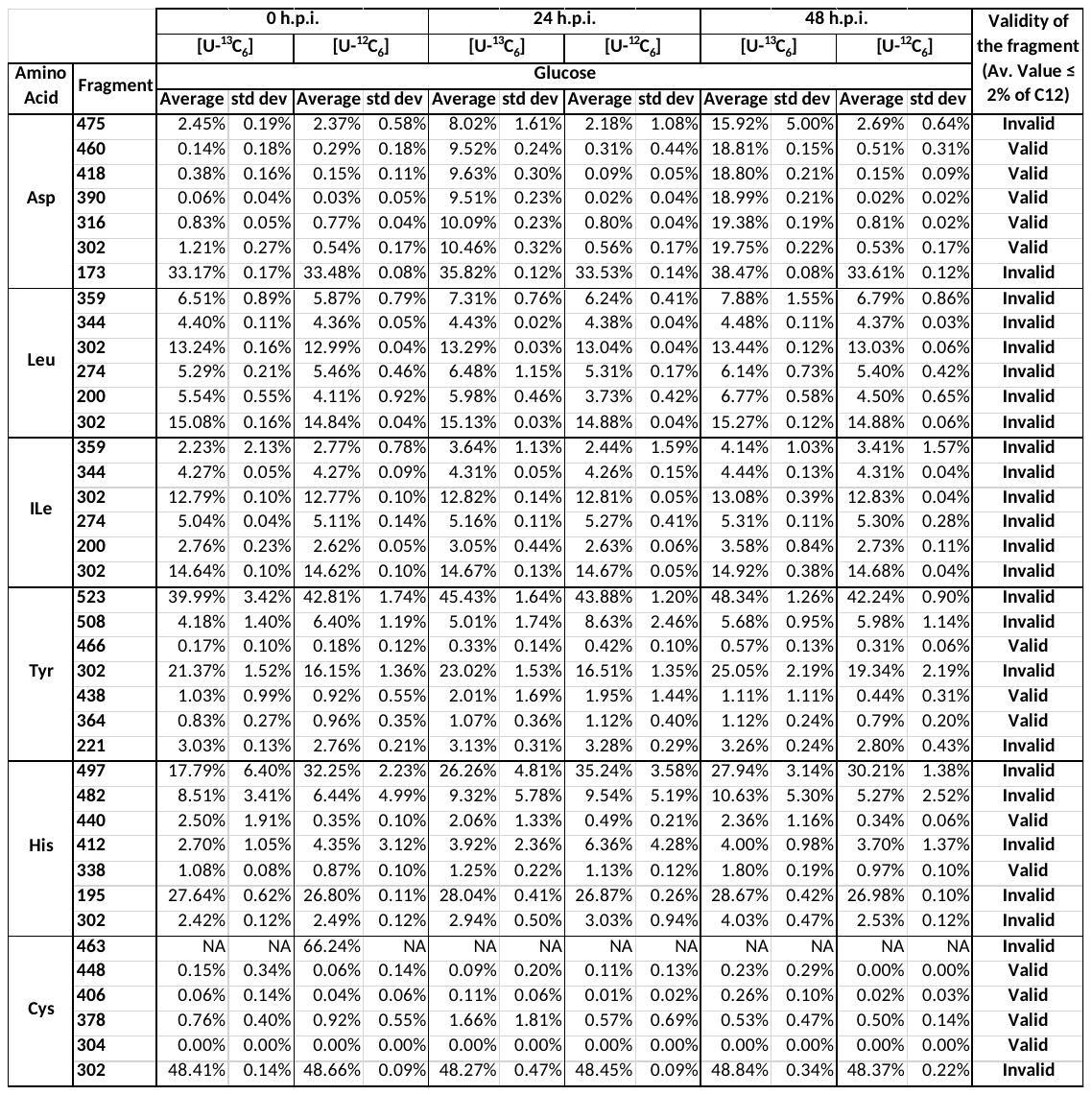

**c. Mtb S4-infected A549 cells**

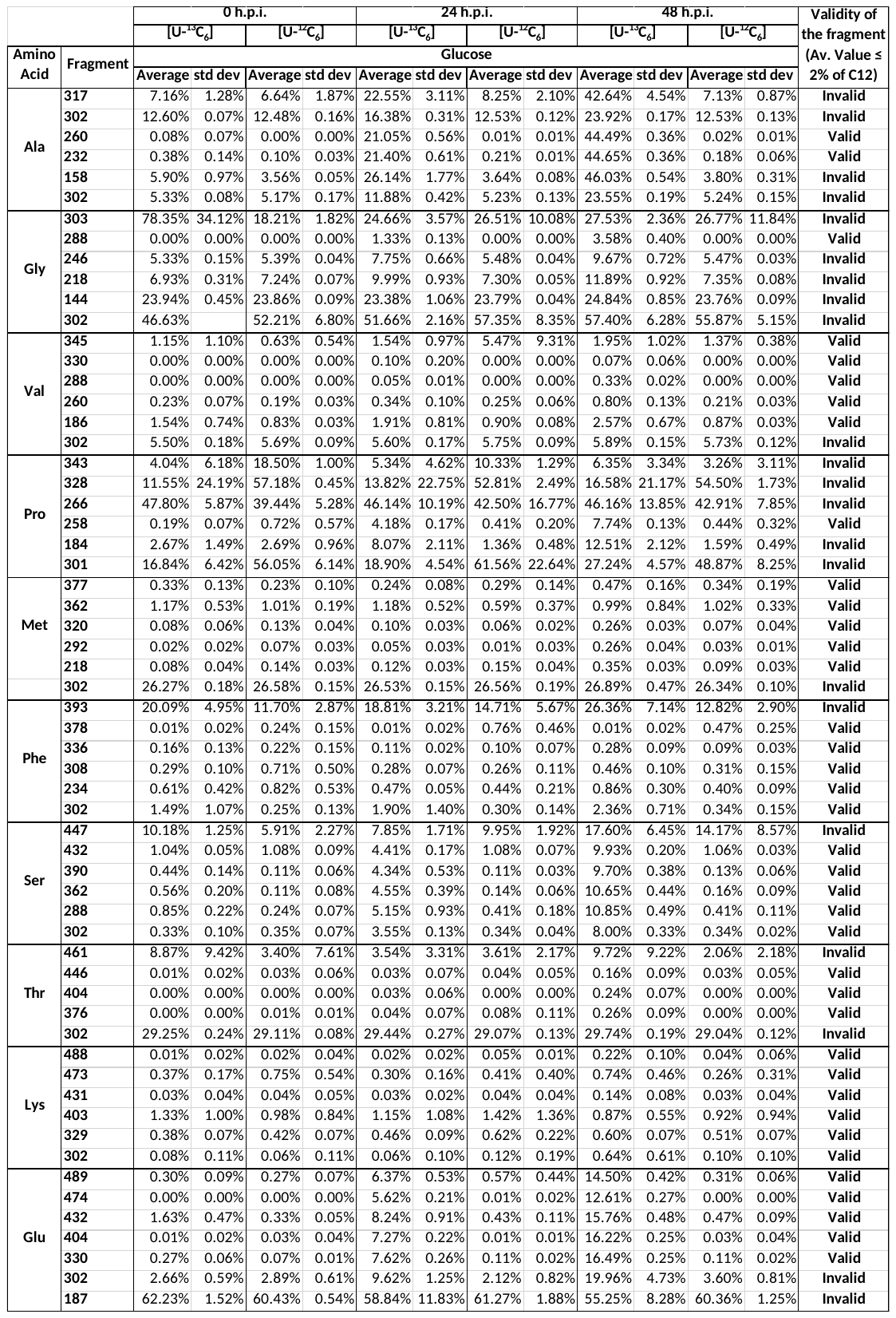

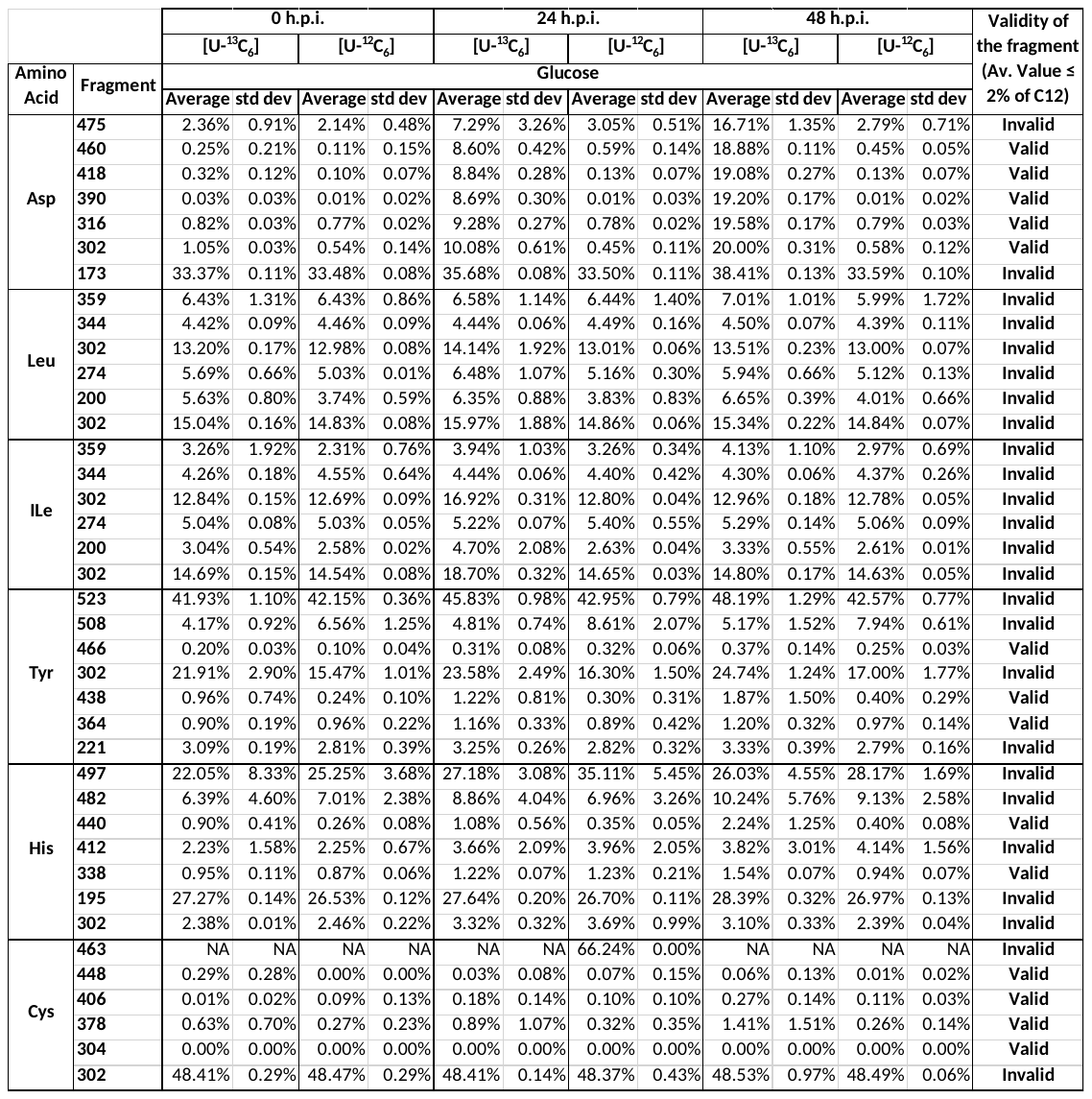

**d. Mtb S5-infected A549 cells**

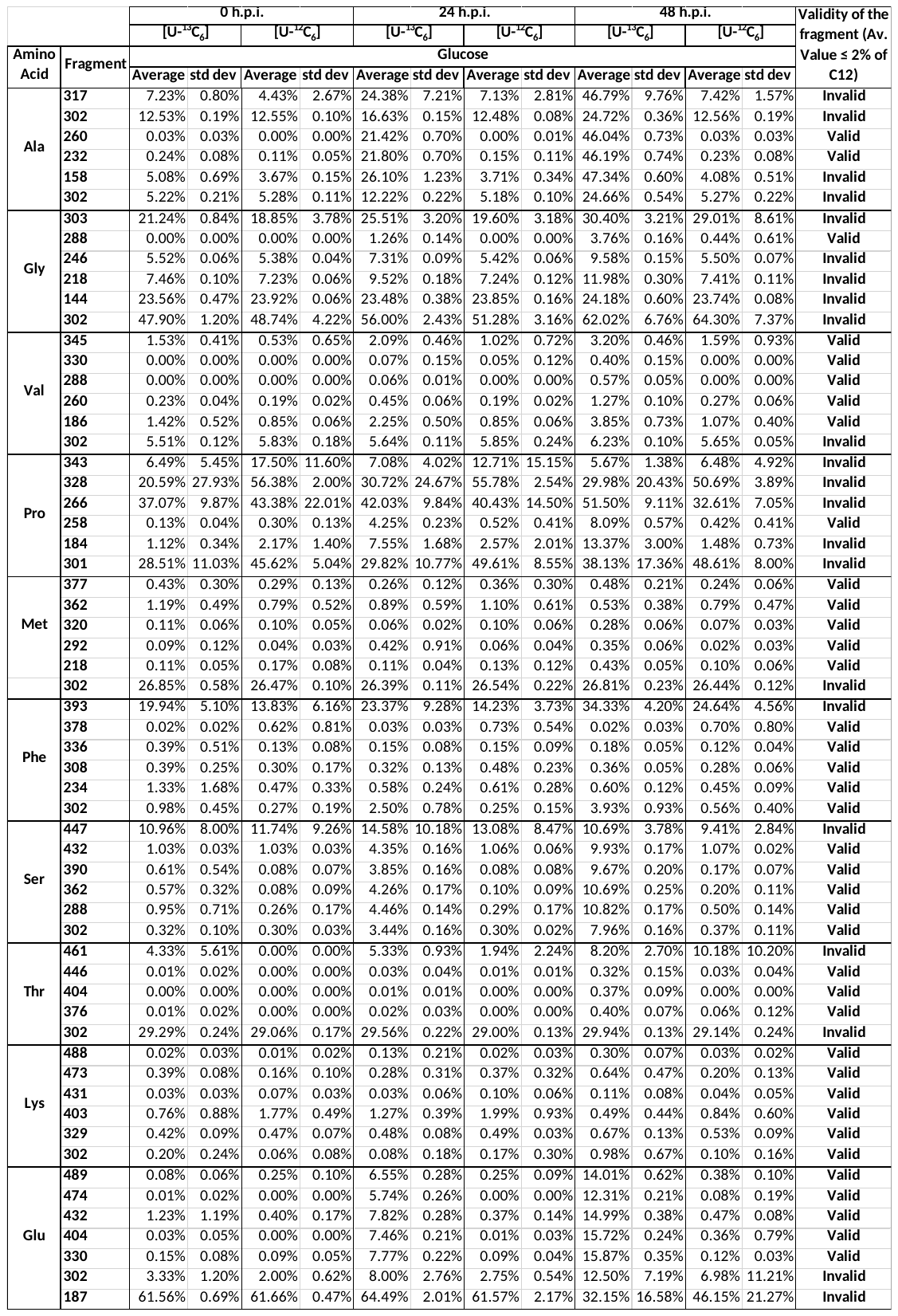

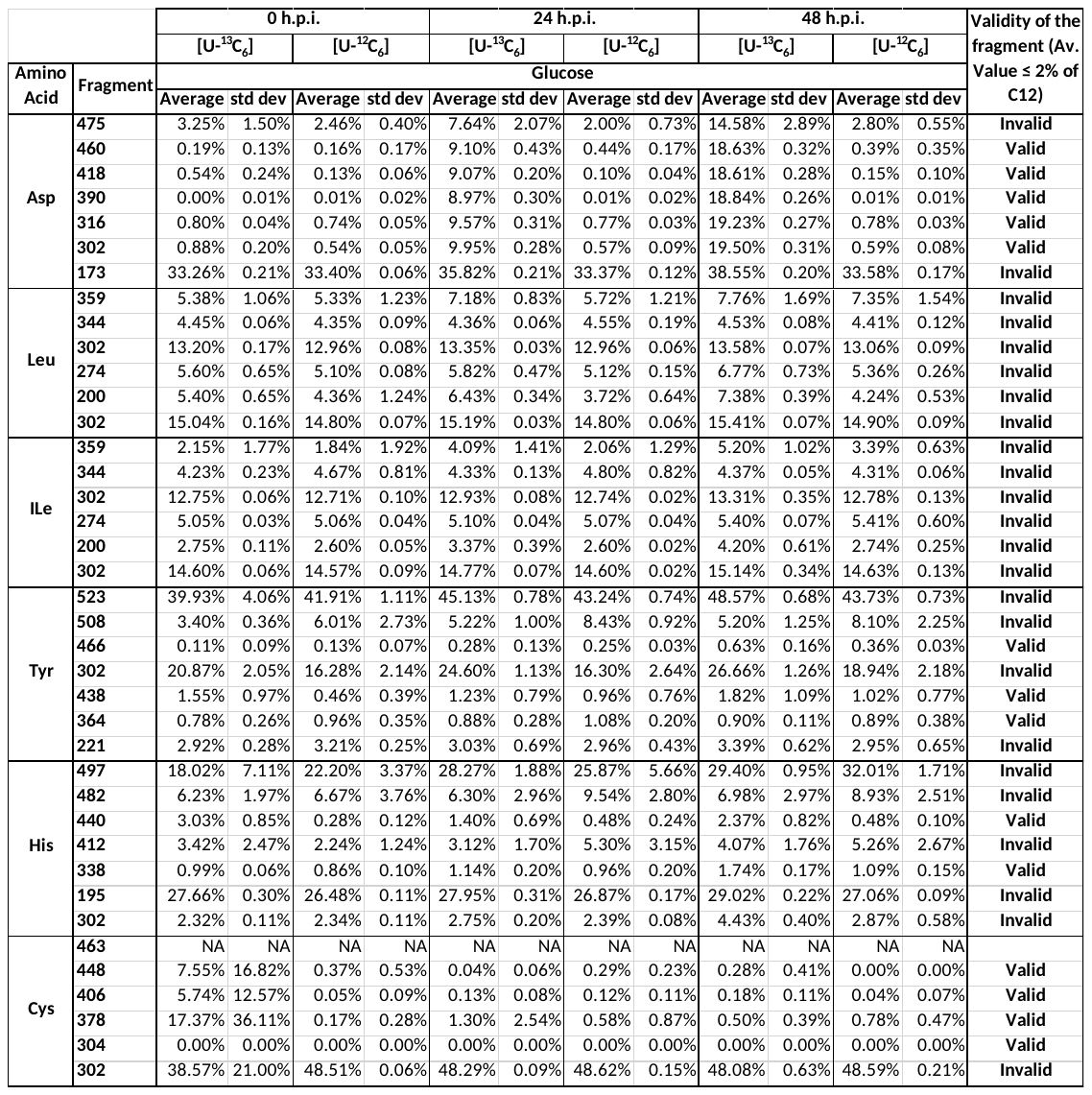

**e. Mtb S6-infected A549 cells**

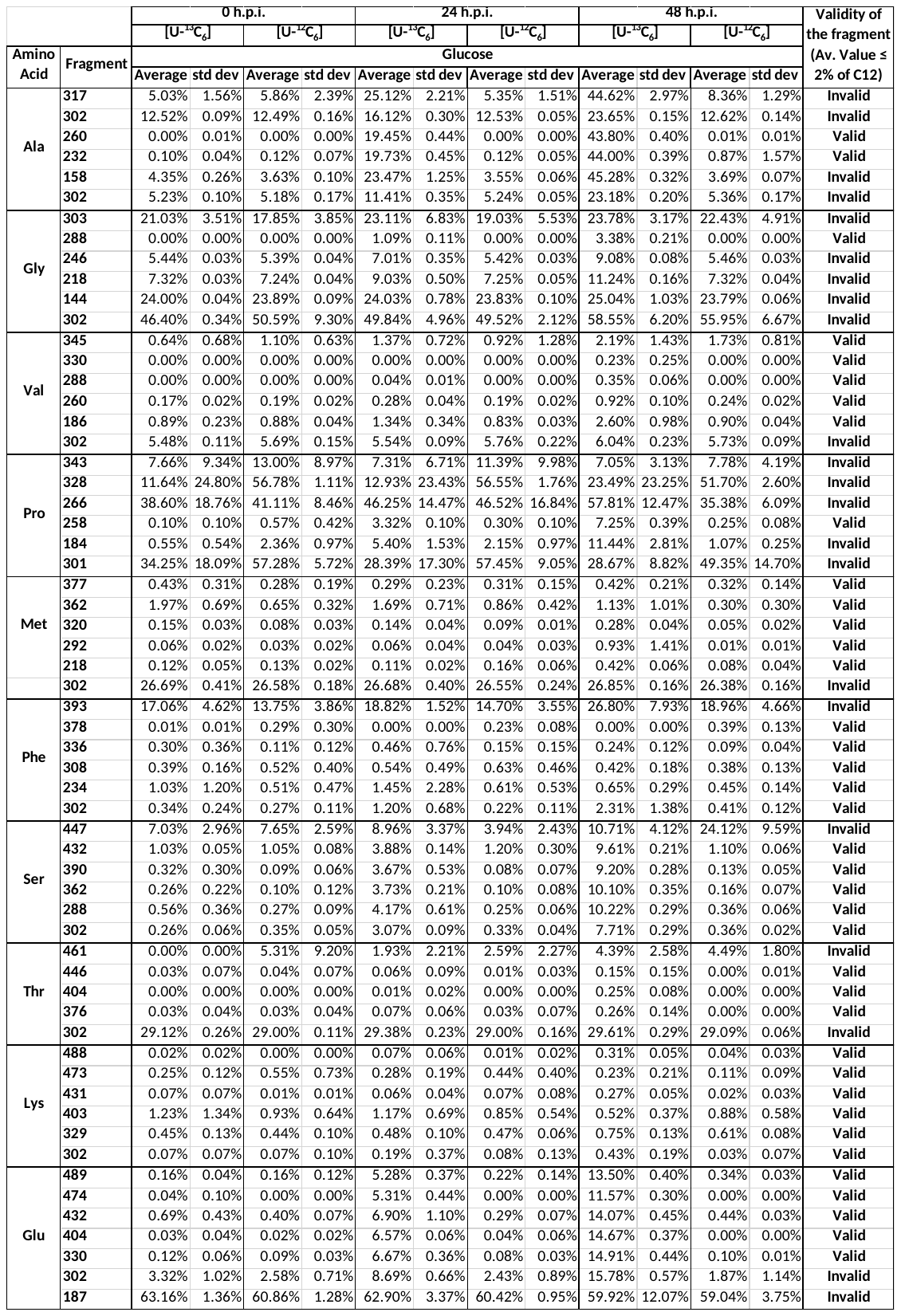

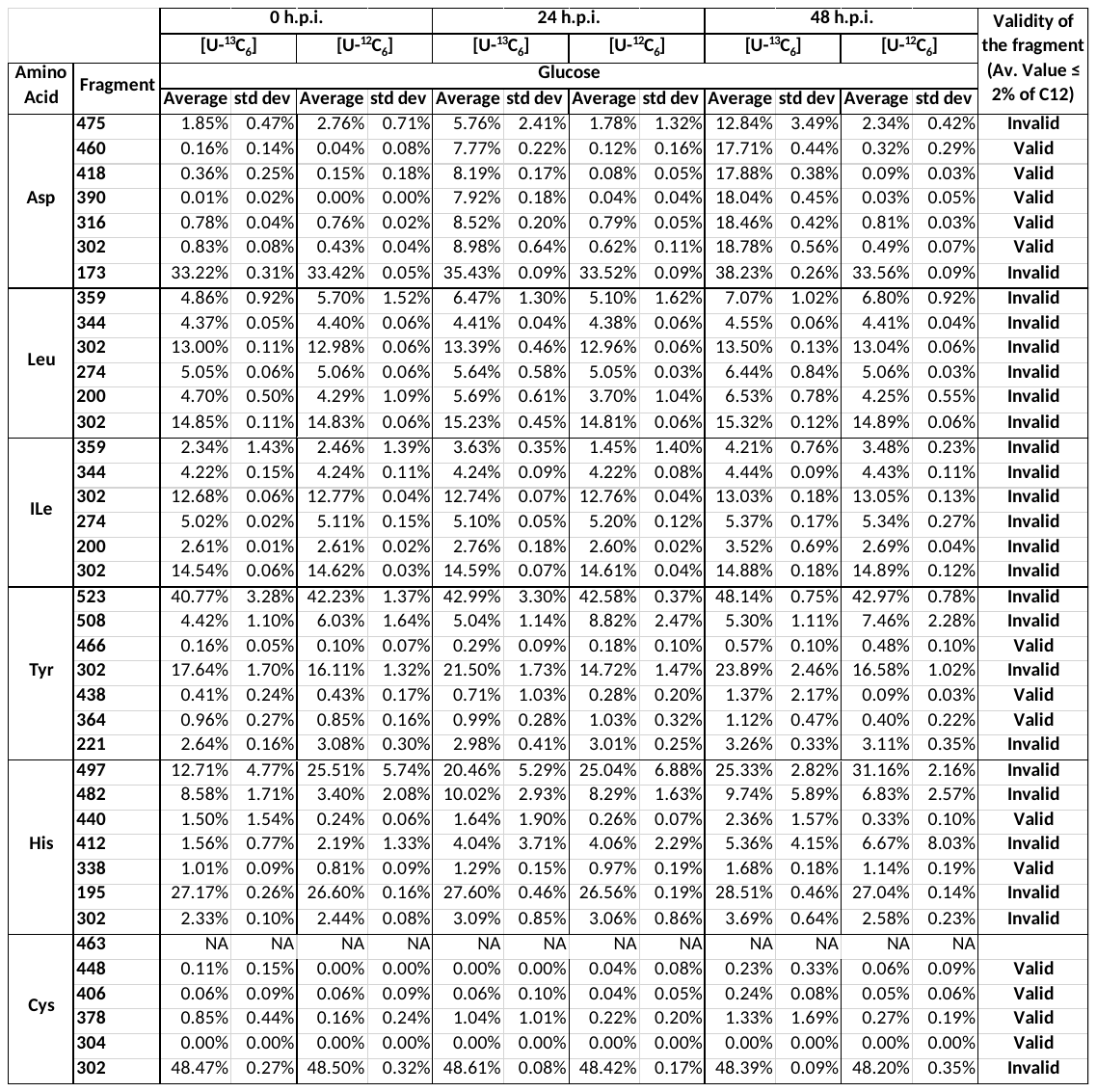

**f. Mtb S11-infected A549 cells**

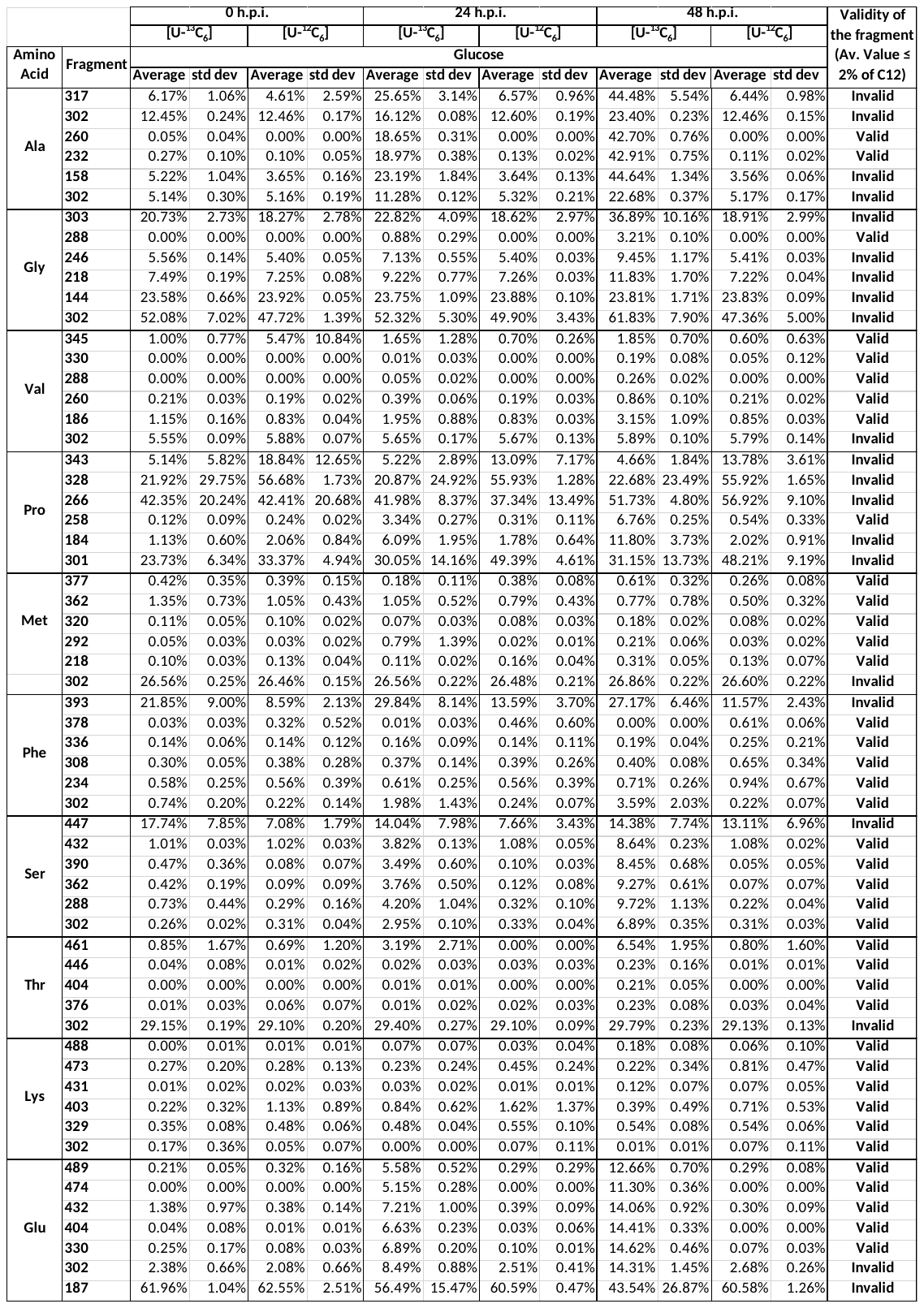

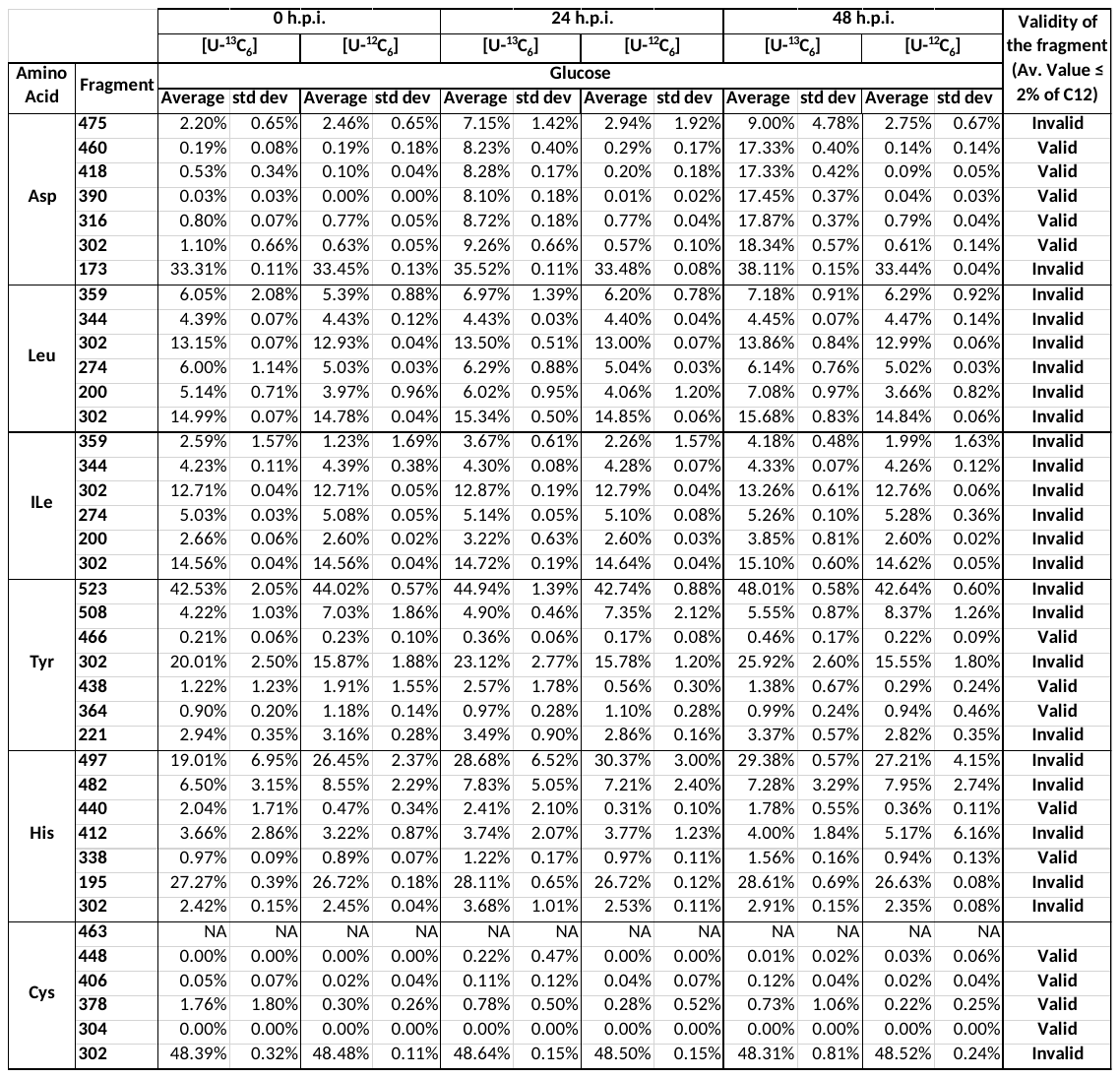

**Supplementary Table 5:** Mass isotopomer distribution (MID) of the proteinogenic amino acid fragments of A549 cells at 48 h.p.i. cultured in DMEM/F12 medium containing either [U-^13^C_6_] or [U-^12^C_6_] glucose and infected with *Mycobacterium tuberculosis* (Mtb) clinical isolates (S4, S5, S6 and S11) and reference laboratory strain H37Rv with uninfected cells as control (replicates=5). Std dev: Standard deviation. Related to Figure 8.

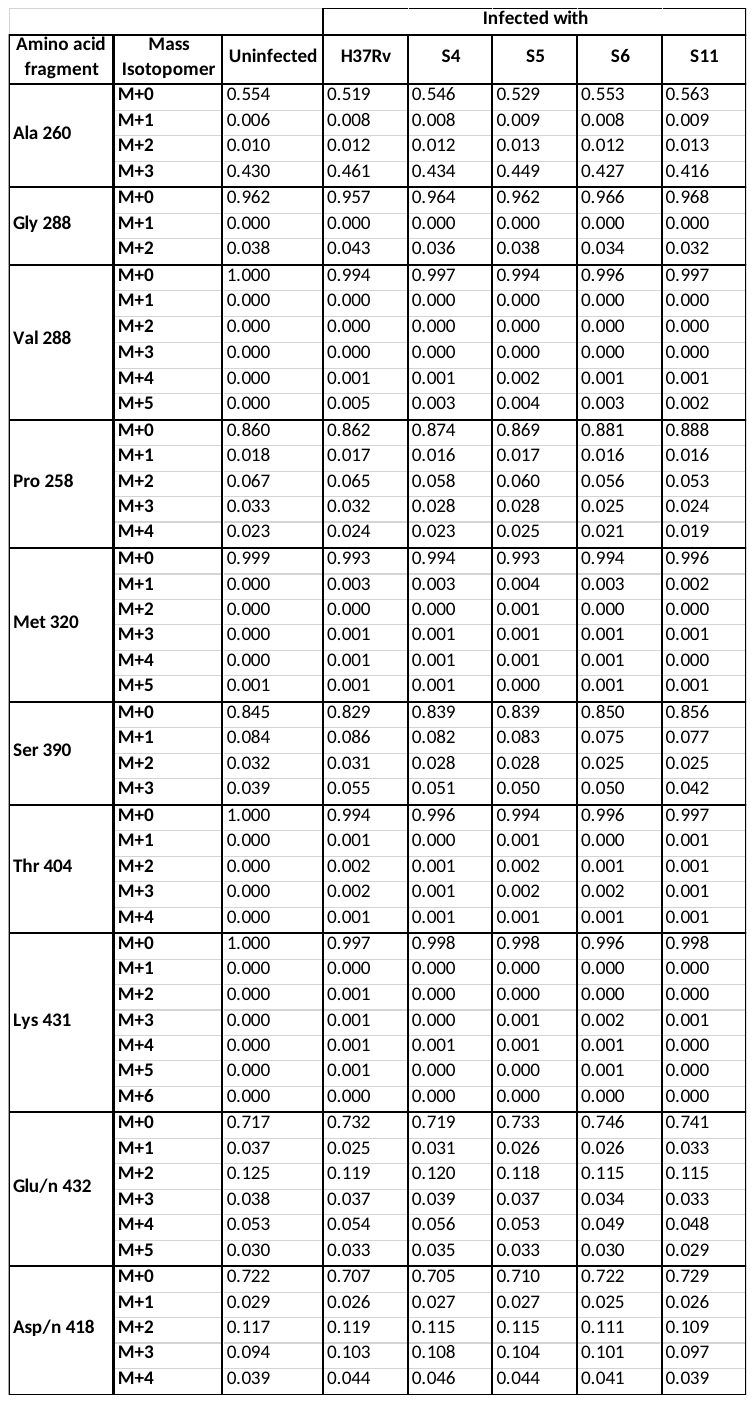

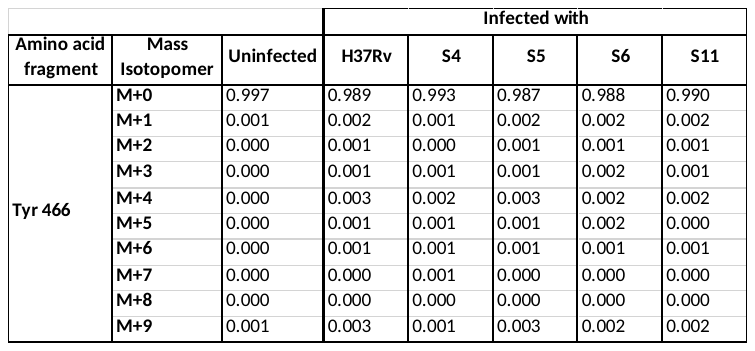

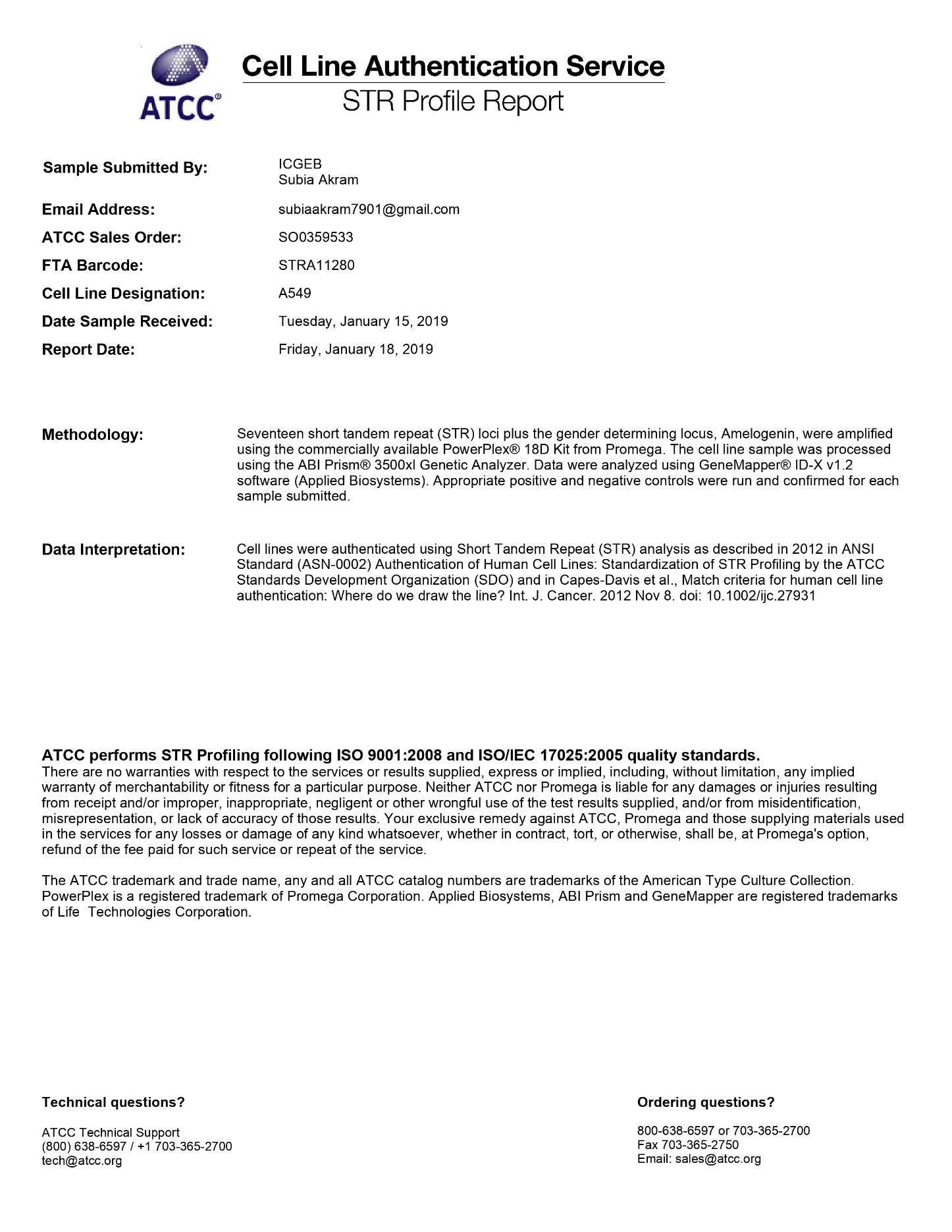

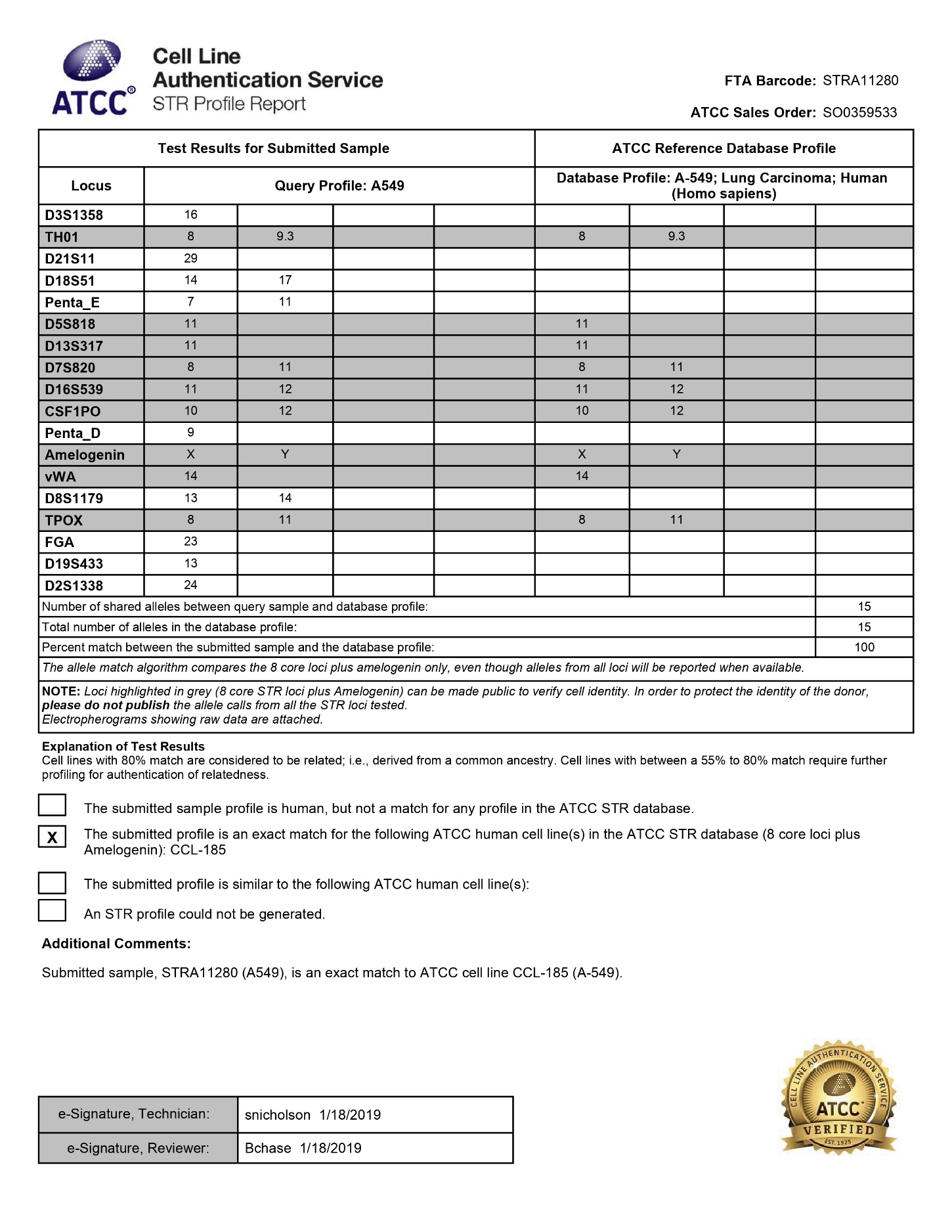
